# Fruitless decommissions regulatory elements to implement cell-type-specific neuronal masculinization

**DOI:** 10.1101/2020.09.04.281345

**Authors:** Margarita V. Brovkina, Rachel Duffié, Abbigayl E. C. Burtis, E. Josephine Clowney

## Abstract

In the fruit fly *Drosophila melanogaster*, male-specific splicing and translation of the Fruitless transcription factor (Fru^M^) alters the presence, anatomy, and/or connectivity of >60 types of central brain neurons that interconnect to generate male-typical behaviors. While the indispensable function of Fru^M^ in sex-specific behavior has been understood for decades, the molecular mechanisms underlying its activity remain unknown. Here, we take a genome-wide, brain-wide approach to identifying regulatory elements whose activity depends on the presence of Fru^M^. We identify 436 high-confidence genomic regions differentially accessible in male *fruitless* neurons, validate candidate regions as bona-fide, differentially regulated enhancers, and describe the particular cell types in which these enhancers are active. We find that individual enhancers are not activated universally but are dedicated to specific *fru*^+^ cell types. Aside from *fru* itself, genes are not dedicated to or common across the *fru* circuit; rather, Fru^M^ appears to masculinize each cell type differently, by tweaking expression of the same effector genes used in other circuits. Finally, we find Fru^M^ motifs enriched among regulatory elements that are open in the female but closed in the male. Together, these results suggest that Fru^M^ acts cell-type-specifically to decommission regulatory elements in male *fruitless* neurons.

## Introduction

In many species, male and female brains generate distinct behavioral repertoires. The ability to compare behavior, brains, neurons, and gene expression across the sexes makes sexually dimorphic behaviors premier models for understanding structure-function relationships in neural circuits. In both vertebrates and invertebrates, master regulators of neuronal sex are induced downstream of the sex determination hierarchy and alter the composition of specific neurons and brain areas (1–4). Circuit changes produced by these transcription factors are complex and heterogeneous, including differences in the numbers of specific types of neurons, their anatomy and connectivity, and their mature physiology (2–7). Collectively, these sex-specific alterations to circuits cause males and females to perform sex-specific innate behaviors. While the causal role of these master regulators in shaping behavior is clear, the transcriptional events through which they do so are opaque.

In insects, neural circuits that regulate mating are masculinized by the action of the *fruitless* transcription factor. Male *Drosophila melanogaster* flies selectively perform courtship displays toward conspecific virgin females, extending and vibrating a single wing to sing a courtship song (8). Fruitless is both necessary and sufficient for masculinization of behavior: While 2-5% of neurons in both male and female brains express *fru* transcript from its sexually dimorphic promoter, fruP1, sex-specific splicing results in functional protein, Fru^M^, only in the male (9–14). Males mutant for *fruitless* exhibit dysregulated courtship behaviors, while females genetically manipulated to produce Fru^M^ protein perform courtship displays to other females (11,12,15–18). Sex-specific morphological differences in >60 classes of *fru*+ neurons have been catalogued (2,3,19), and many of these dimorphic populations are implicated in the regulation and performance of male mating behaviors (8).

The *fru*+ neurons are found in every part of the nervous system, including sensory structures, the central brain, and motor output regions (9,10). Most subclasses derive from distinct neuroblasts and are only a small proportion of the cells born from each neuroblast (3). The sexual differentiation of different types of *fruitless* neurons produces changes to male and female cell number, anatomy, connectivity, function, or a combination of these (2,3,19–22). Work from many labs over the last 15 years has suggested that *fruitless* neurons throughout the nervous system preferentially interconnect to form a modular circuit dedicated to sex-specific behaviors (20,21,23,24).

Fru^M^ is a sequence-specific DNA binding protein that likely turns on and off the transcription of specific gene targets (11,12). Previous studies have identified a handful of Fru^M^ effectors but have not addressed broader logical principles about what it means to be a male neuron (25–27): Does Fru^M^ act on the same targets across different *fruitless* cells? Are the genes it regulates dedicated to the mating circuit? Does it stimulate or repress transcription? Ultimately, defining the transcriptional role for Fru^M^ in masculinizing this circuit will allow us to ask how differences in overall circuit architecture and behavior emerge from independent gene regulatory events in its cellular constituents (28).

To increase resolution for identifying candidate enhancers and repressors directly or indirectly regulated by Fru^M^, we performed the Assay for Transposase-Accessible Chromatin (ATAC-seq) (Buenrostro et al 2015) on FAC-sorted *fru*^+^ and *fru*^-^ neurons from male and female brains. Using this exquisitely sensitive method, we identified 436 genomic elements differentially accessible in the presence of Fru^M^. To measure the gene regulatory activity of these elements and to define the specific subpopulations of *fru*^+^ cell types in which they are capable of regulating gene expression, we analyzed the ability of matched genomic fragments to drive reporter expression across the brain. The combination of these genome-wide and brain-wide approaches allows us to define cell-type-specific enhancers dependent on Fru^M^ and the logic of Fru^M^ action across different populations of *fru*^+^ cells. We find that individual regulatory elements differentially accessible in the presence of Fru^M^ are each used in only a small population of *fru*^+^ neurons, suggesting that each subpopulation of *fru*^+^ cells has a distinct set of Fru^M^ effectors. We therefore conclude that Fru^M^ acts as a “switch gene” (28): It flags cells as male and interacts differently with the cell-type-specific transcriptional milieus of different cell types to induce diverse masculinizing adjustments. While regulatory elements may be dedicated to specific Fru^M^ populations, the genes they regulate are shared with other circuits. Finally, we identify differentially accessible genomic regions with strong Fru^M^ motifs; these are enriched in regions specifically closed in the presence of Fru^M^ protein, suggesting that Fru^M^ acts to decommission its direct targets.

## Results

To determine how *fruitless*-expressing neurons in male flies are differentially patterned to allow males to perform distinct behaviors from females, we sought to identify genetic elements whose regulatory state correlated with the presence of Fru^M^ protein. As chromatin accessibility often correlates with the activity of regulatory elements, we used ATAC-seq, a method for identifying open chromatin regions genome-wide (29). *fruitless* is expressed across much of the life cycle: expression begins in late larvae, peaks in mid-pupae, and continues robustly in the adult (13). Fru^M^-dependent sexual dimorphisms, including in developmental apoptosis, neuronal arbor patterning, and functional properties, encompass differences that likely arise at each of these time points (6,20,30–33). We chose to complete our initial analysis at the adult stage, as comprehensive knowledge of the repertoire of adult *fruitless* neurons allows us to identify the neurons in which individual regulatory dimorphisms occur.

We used expression of GFP under control of fruP1-Gal4 (10) to report transcription from the fruP1 promoter, and then FAC-sorted and analyzed four populations of cells: *fru*^+^ and *fru*^-^ cells from male and female (Figure 1A-C). These four populations allow us to define transcriptional differences related to sex, *fru* transcriptional status, and Fru^M^ protein status.

**Figure 1.**
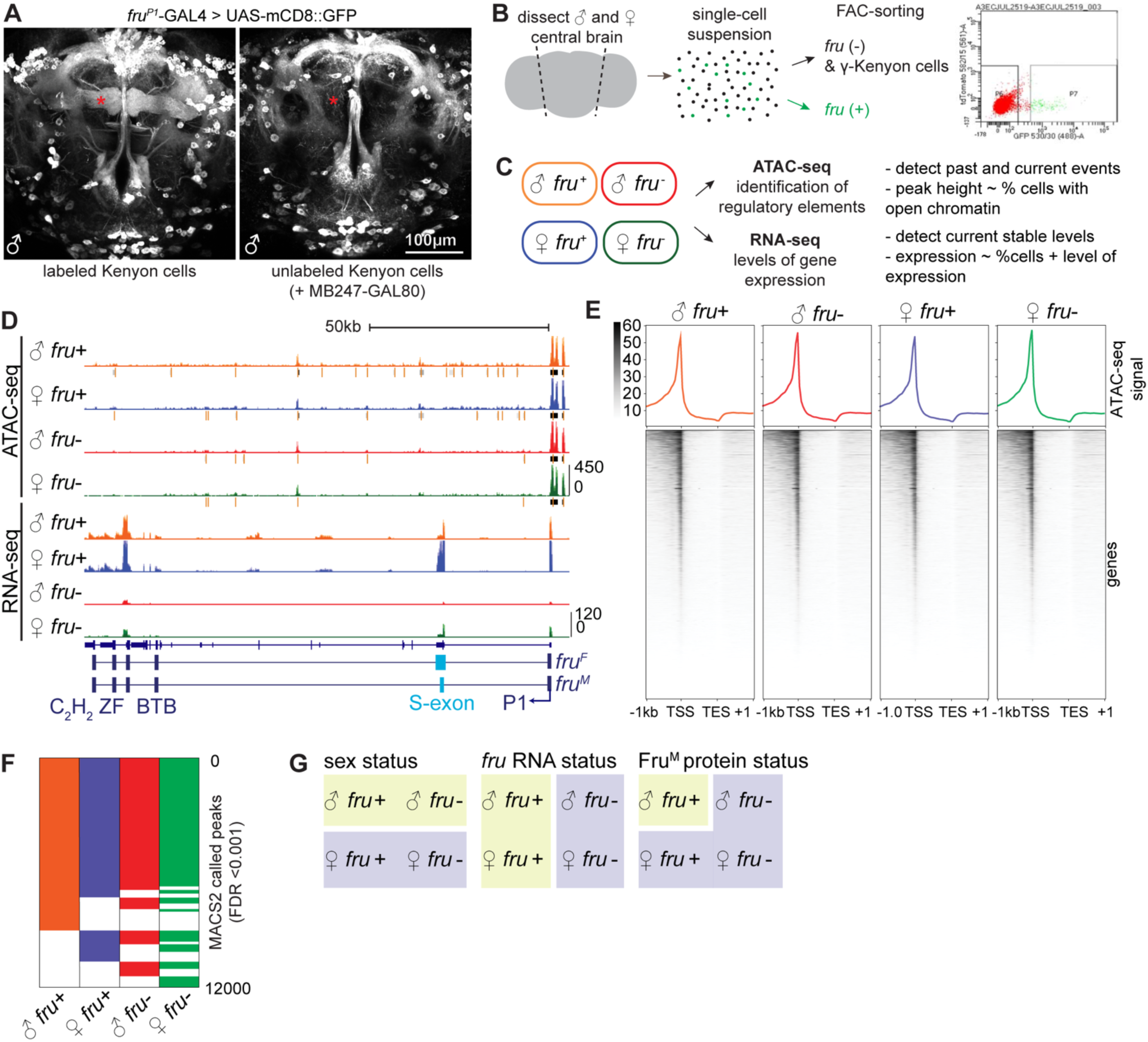
Genomic profiling of a core population of *fruitless* neurons. A. 2-photon maximum intensity projection of fruP1-GAL4 driving mCD8-GFP in the adult male central brain, shown with and without MB247-Gal80 masking GFP signal in mushroom body Kenyon cells. Here and throughout, brain images show anterior view. B. Scheme of dissection and sorting; example FACS plot is shown. C. Scheme of datasets collected. D. UCSC genome browser screenshot of the *fruitless* locus. *fruitless* transcript structure is schematized below. Here and throughout, each signal track shows a transparent overlay of two independent biological replicates. Gray bars under ATAC signal indicate MACS2-called peaks at FDR threshold <0.001, with orange bar indicating peak summit. E. Signal heatmap of ATAC signal (average of two independent biological replicates) summarized over all annotated genes. F. Binary heatmap of genomic regions which contain a peak of open chromatin (MACS2 FDR <0.001). G. Axes that separate the four analyzed cell populations.

**Figure S1.**
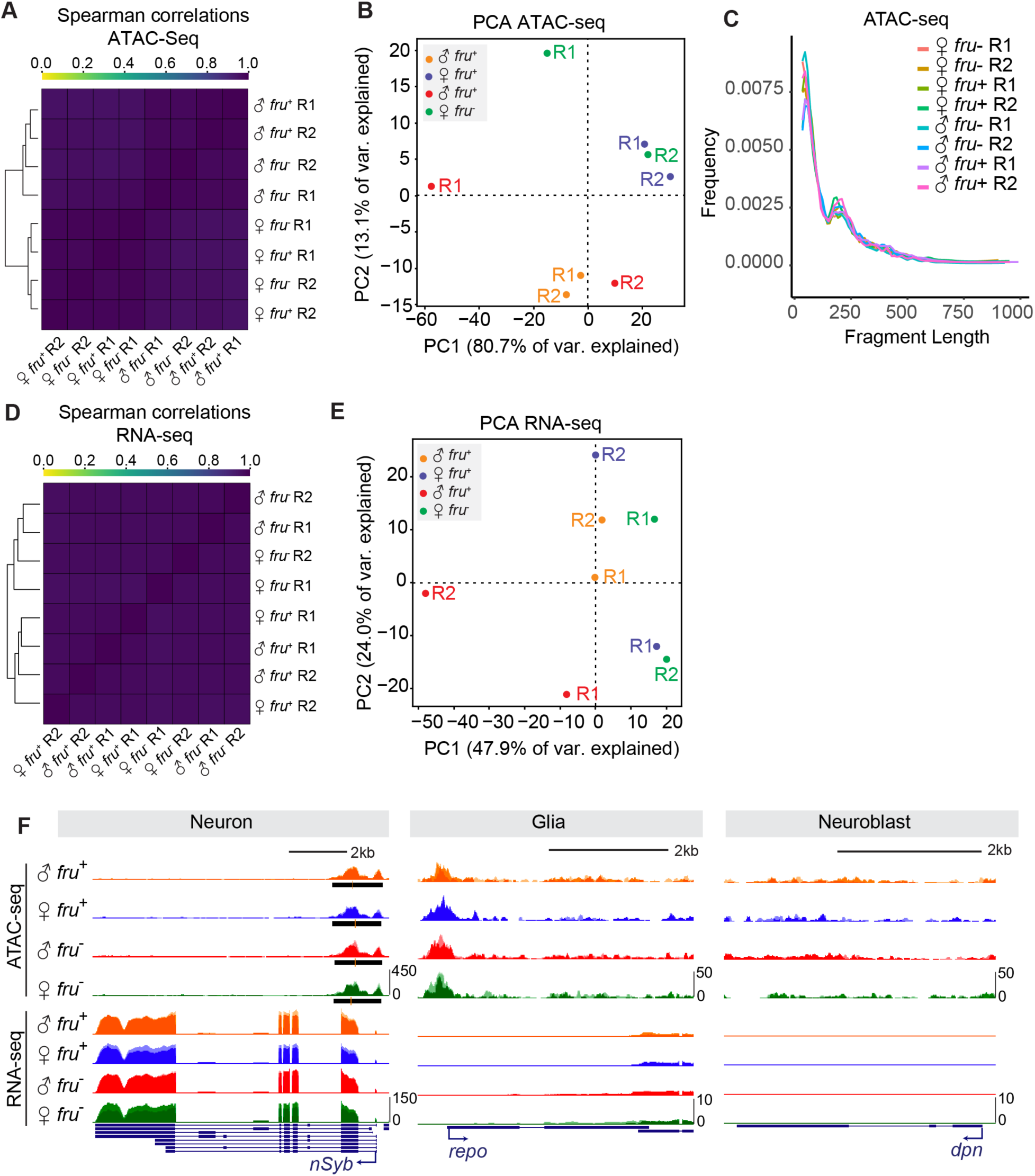
Quality control metrics for ATAC and RNA sequencing. A. Heatmap of Spearman correlations of uniquely aligned, deduplicated reads from ATAC-seq libraries. B. PCA analysis of uniquely aligned, deduplicated reads from ATAC-seq libraries. PC1 reflects read depth. C. Fragment length of ATAC-seq libraries D. Heatmap of Spearman correlations of uniquely aligned, deduplicated reads from RNA-seq libraries. E. PCA analysis of uniquely aligned, deduplicated reads from RNA-seq libraries F. UCSC genome browser screenshots of ATAC-seq and RNA-seq signal across neural (*nSyb*), glial (*repo*), and neuroblast (*dpn*) specific genes.

*fruitless* neurons in the central brain are morphologically diverse and derive from >60 neuroblasts (2,3,19). To enrich for cell populations of interest, from the central brain, we removed optic lobes and the ventral nerve chord. In addition, the largest population of *fruitless* neurons is γ Kenyon cells. Kenyon cells are required for olfactory learning, including courtship learning; however, Kenyon cells are not required for core courtship programs (34). To prevent these numerous cells from dominating our analyses, we used MB247-Gal80 to remove reporter expression in γ Kenyon cells and therefore sorted them into the *fru*^-^ populations (Figure 1A-B); we estimate that γ KCs comprise 3% of our *fru*^-^ libraries. For each of two biological replicates, we sorted GFP^+^ cells, and matched numbers of GFP^-^ cells, from >20 male and female brains; this yielded 6,000-10,000 cells per sample. GFP^+^ cells were 1.8-3.6% of cells in female, and 3.6-5% of cells from male, in line with previous estimates of *fru*^+^, non-Kenyon cells from the central brain (10,13). Samples were then subjected to ATAC-seq (two replicates) or RNA-seq (two replicates) (Figure 1C, D).

*fruitless* is transcribed from several promoters: The fruP1 promoter drives expression of the sexually dimorphic transcripts, including the male transcript coding for Fru^M^, while three downstream promoters drive expression of the Fru^COM^ protein, which is not dimorphic and which is required in both sexes for embryogenesis (12,14). We do not observe fru^COM^ transcripts in adult neurons (Figure 1D), and previous analyses have found that Fru^COM^ protein is not present after pupation (13), thus ensuring that our analyses are restricted to differences that arise from the sexually dimorphic Fru^M^ protein.

We captured strong (∼4.8 fold) enrichment of *fruitless* transcript in the *fru*^*+*^ RNA-Seq libraries. The small amount of *fru* mRNA signal in the *fru*^*-*^ libraries is expected to derive from the *fruitless* mRNA expressed in γ Kenyon cells, which we sorted into the *fru*^*-*^ pool. We observe *fru* mRNA in both male and female cells, with the expected sexually dimorphic splicing of the S exon clearly visible. While close to 100% of the *fru* transcript in female cells matches the expected female splicing variant, a small subset of transcripts in the male have the female exon. In addition to sexually dimorphic splicing, *fru* transcripts are alternatively spliced at the 3’ end, yielding several possible DNA binding domains (12). Protein products of these isoforms are designated Fru^MA^, Fru^MB^, and Fru^MC^, and previous results have shown that expression of these three isoforms is largely overlapping, such that most *fruitless* neurons contain all three (30). We observe all three 3’ splice isoforms in our libraries. QC metrics are presented in Figure S1A-F.

For ATAC-seq, we isolated nuclei and subjected them to TN5 transposition, DNA purification, library preparation, and paired-end sequencing (29), with modifications made for low input cell numbers (35). After mapping reads to the *D. mel* genome (dm6) and removing duplicates, we obtained 9-16 million distinct reads per sample. The eight libraries showed strong Spearman correlation overall, as expected given their common source (Figure S1A). Male and female libraries clustered separately, and male samples clustered according to *fru* status, while female samples, all of which lack Fru^M^ protein, intermingled (Figure S1A). Likely due to the high overall correlations, principle component analysis was dominated by read depth (Figure S1B). We aggregated ATAC-seq signal across genes and found strong enrichment at promoters, weaker enrichment 3’ of genes, and depletion of signal from transcribed regions (Figure 1E). We then used MACS2 to call peaks in each of the four cell types, yielding >11,000 peaks genome-wide for each sample at FDR<0.001 (36). ∼60% of peaks were called universally across the four sample types, and the remaining 40% were condition-specific (Figure 1F). While we observe many intronic peaks in our dataset, these peaks occupy diverse positions in TSS-anchored gene models and thus do not show strong aggregate signal (Figure 1E). Comparisons across these four cell types allow us to investigate three axes of neuronal difference: sex, *fru* transcript status, and Fru^M^ protein status (Figure 1G).

### Sexually dimorphic, Fru^M^-independent peaks reflect dosage compensation

Sexually dimorphic splicing of fruP1 transcripts represents a late, tissue-specific event in the sex differentiation hierarchy (11,12). In order to test the validity of our data, we first assessed whether we could detect hallmarks of X chromosome dosage compensation in adult neurons, which is expected to be sexually dimorphic but Fru^M^ independent (Figure 2A). In *Drosophila melanogaster*, the dosage compensation complex (DCC) binds to ∼700 regions on the male X to upregulate gene expression chromosome-wide (Figure 2A) (37,38). Two X-linked lncRNAs that help to target the DCC to the male X, *roX1* and *roX2*, were expressed only in our male samples and expressed similarly in *fru*^*+*^ and *fru*^*-*^ cells, as expected (Figure 2B) (39). We also observe male-specific ATAC-seq signals at the *roX1* and *roX2* loci (Figure 2B).

**Figure 2.**
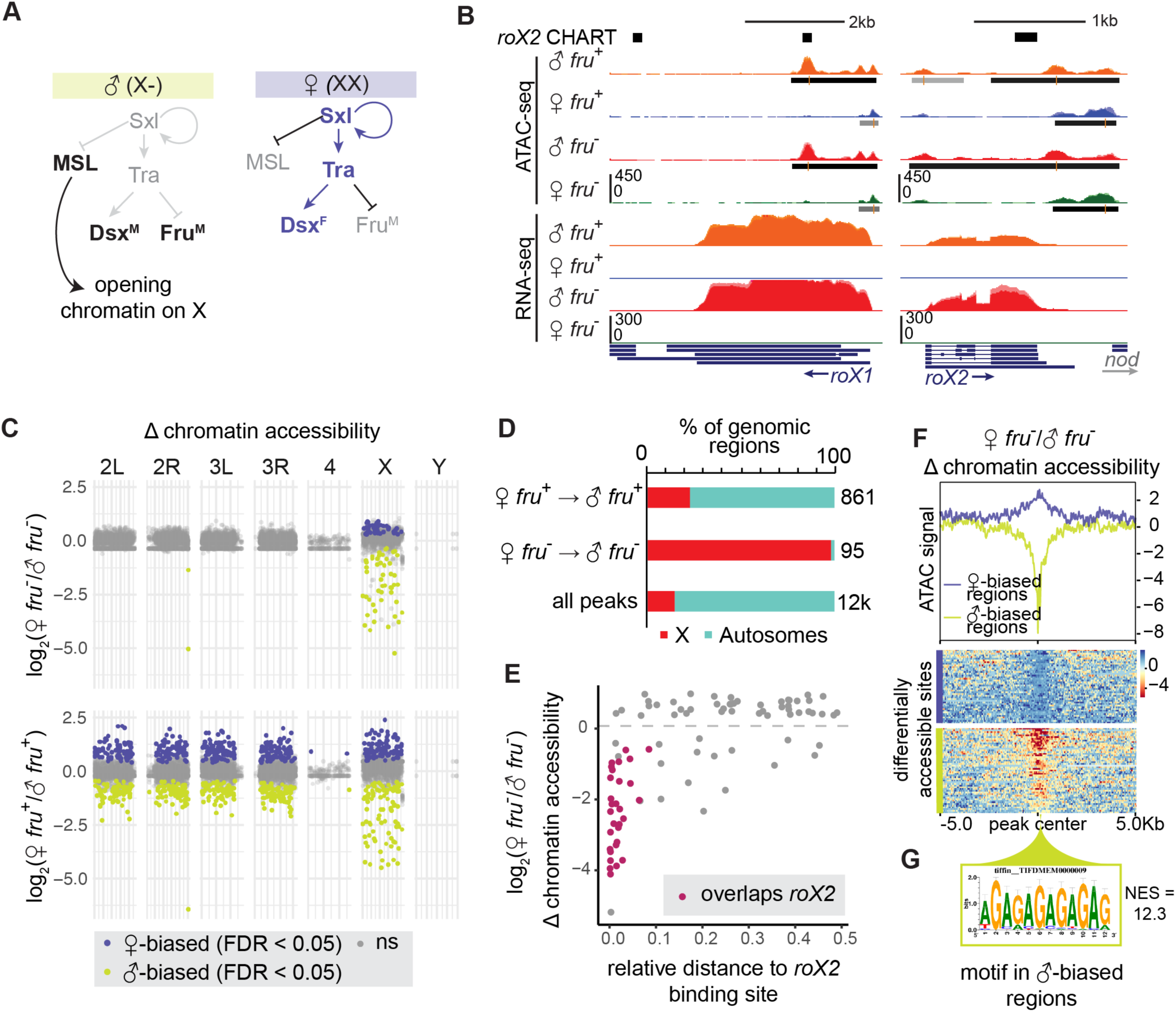
Sexually dimorphic, Fru^M^-independent peaks reflect dosage compensation. A. Schematic of dosage compensation in *D. mel*. B. UCSC genome browser screenshot of ATAC-seq and RNA-seq signal at canonical dosage compensation genes *lncRNA:roX1* and *lncRNA:roX2. roX2* CHART (black bars) shows known tethering sites of *lncRNA:roX2* to chromatin (38). C. Manhattan plot of log_2_(female/male) chromatin accessibility in *fru*^+^ and *fru*^-^ neurons across chromosomes. Points represent genomic regions. Vertical axes show the relative change in accessibility of a region from DiffBind output. Horizontal axes represent scaled chromosomal locations. Regions with no change in accessibility between samples (FDR > 0.05) are plotted in gray. Regions with male-biased accessibility (chartreuse) appear as negative fold change, and regions with female-biased accessibility (purple) appear as positive fold change. D. Summary of X versus autosome distribution of differentially accessible genomic regions (DiffBind, FDR<0.05) versus all peaks (MACS2, FDR<0.001). E. Relative distance of regions specifically accesible in female versus male *fru*^-^ neurons to *lncRNA:roX2* binding sites identified by CHART (38). Sex-biased regions which directly overlap a roX2 tethering site are highlighted in magenta. F. Signal heatmap of log2 fold change (log2FC) in ATAC-seq coverage between female *fru*^-^ and male *fru*^-^ neurons. Line plot shows mean log2FC in signal across reference-anchored regions. Signal heatmaps below are split into regions with female-biased (purple) or male-biased (chartreuse) accessibility. G. I-cisTarget analysis of male-biased regions shows enrichment of a GAGA motif matching the known CES motif for the dosage compensation machinery. The normalized enrichment score of the motif is 12.3.

**Figure S2.**
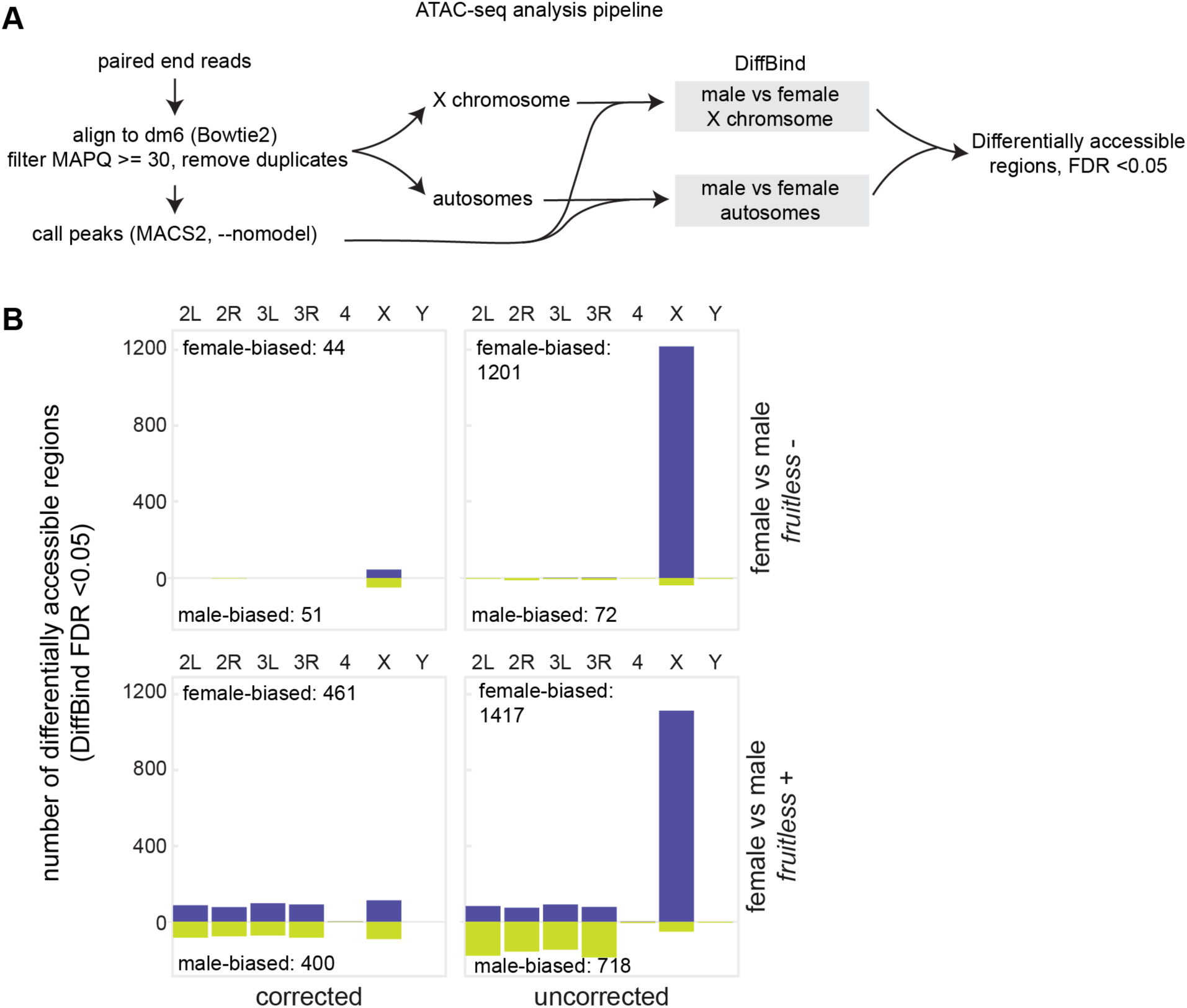
Novel analysis pipeline to correct for X:A autosome ratio in *D. mel*. A. Flowchart of computational pipeline used to identify differentially accessible regions between male and female samples using ATAC-seq. B. Barplots displaying number of differentially accessible regions using DiffBind FDR <0.05 as a cutoff. Corrected regions are produced by the pipeline in A, while uncorrected regions are produced without splitting of the X chromosome and autosomes before running DiffBind. Colors correspond to Fig 2C, where purple are regions that have female-biased accessibility, and chartreuse represents regions with male biased accessibility.

To define differentially accessible peaks across our sex-specific datasets, we first developed methods to correct for the different numbers of X chromosomes in male and female. When the X chromosome and autosomes were analyzed together using common pipelines, most differential peaks genome-wide were found to be female-biased and from the X chromosome, presumably due to the two-fold difference in genetic material from female versus male X. As the X chromosome comprises a large proportion of the fly genome, the bias this induced in our analysis was strong: the X chromosome constituted ∼20% of our total ATAC-seq reads in female flies, and only 14% in male flies. To correct for this disparity, we separated X chromosome and autosome reads in each dataset (Figure S2A). We then used DiffBind to call differential peaks on autosomes alone, and on the X chromosome alone (40,41). This method increased our sensitivity to detect differential peaks on the autosomes between the two sexes and allowed us to predict true accessibility differences on the X chromosome (Figure S2B).

For all MACS2-called peaks genome-wide, we calculated the change in chromatin accessibility between *fru*^*-*^ male and female samples, and between *fru*^*+*^ male and female samples (Figure 2C, D). The genomic distribution of differential peaks was strikingly different between the *fru*^*-*^ samples and between the *fru*^*+*^ samples. In *fru*^*-*^ samples, 93 of 95 differential peaks were located on the X chromosome, suggesting that the main differences in chromatin accessibility between these samples derived from the process of dosage compensation itself. In contrast, nearly 80% of differentially accessible regions between male and female *fru*^*+*^ neurons were on autosomes (Figure 2C, D). Together, these patterns validate our ability to identify Fru^M^-dependent regulatory events in ATAC-seq data, and allow us to subtract signatures of dosage compensation from our analysis.

On the X chromosome, we observed ∼44 regions that were strongly male-biased, similar to signals in *rox1* and *rox2* (Figure 2B, C, E, F). These peaks, which we predicted represented DCC binding sites, were common to both *fru*^*+*^ and *fru*^*-*^ samples. To test this, we took advantage of known male-specific DCC binding profiles mapped by analysis of *rox2* chromatin binding by CHART (capture hybridization analysis of RNA targets) (38). Indeed, male-biased peaks were relatively closer to DCC binding sites identified by *roX2* CHART than were female-biased peaks, and >50% overlapped DCC binding sites (Figure 2E). Using i-cisTarget, we found sharp enrichment of the MSL recognition element (MRE) in these sequences (Figure 2F, G) (42).

### Transcriptional regulation of *fruP1*

We next characterized accessibility differences between the *fru*^+^ versus *fru*^-^ datasets. We found just 22 regions genome-wide whose accessibility was common to the two *fru*^+^ or two *fru*^-^ datasets, consistent with a model where *fruitless* neurons are heterogeneous in the absence of Fru^M^ protein production (Figure S3A, B). To ask whether these morphologically and lineally distinct neurons activate *fruitless* transcription through common or distinct mechanisms, we next turned our attention to the *fru* locus itself. *fruitless* has a high intron:exon ratio, which has been suggested to correlate with diversification of cis-regulatory elements and expression patterns (43,44). The *fruP1* promoter was accessible regardless of transcriptional status (Figure 1D); hierarchical control of promoter opening versus transcription has been observed in other neural systems (45). There was an enrichment of called peaks in the locus in the two *fru*^+^ datasets, particularly in the first two introns downstream of *fruP1* (Figure 3A). There also appeared to be pervasive enrichment of reads across the locus in the *fru*^+^ datasets. We reasoned that if each subpopulation of *fru* neurons uses a different enhancer within the *fru* locus to activate transcription from *fruP1*, averaging of these accessibility signals across a variety of *fru*^+^ cell types could lead to the observed pervasive opening. To quantify accessibility across the 120kb locus, we measured coverage, i.e. number of reads per base pair. We observed up to 4-fold enrichment in reads across the locus as a whole in *fru*^+^ samples compared to *fru*^-^ samples (Figure 3B). We re-mapped histone-mark ChIP from adult brain neurons (46) and found that this region was also enriched for H3K27Ac, a mark of active enhancers (Figure 3A).

**Figure 3.**
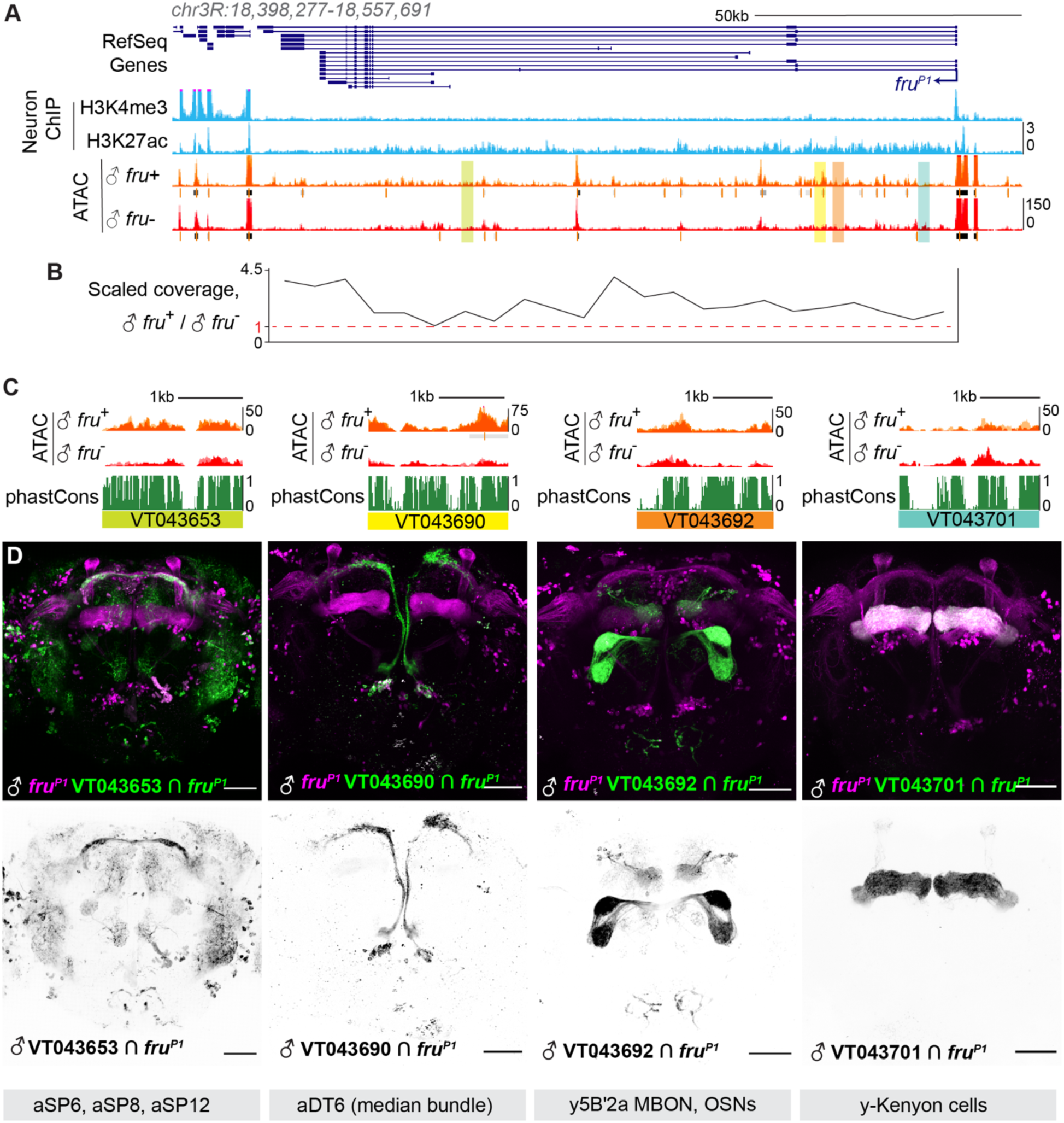
Diverse regions of the *fruitless* locus act as enhancers in subsets of *fru*^+^ neurons. A. UCSC genome browser screenshot of the *fruitless* locus (dm6 assembly). Blue signal tracks represent INTACT histone ChIP data for R57C01(*Nsyb*)-labeled neurons in the adult head (46). The P1 promoter of *fru* shows enrichment of H3K4me3, a promoter mark, while the whole gene body shows signal for H3K27ac, an enhancer mark, compared to low signal upstream of fruP1. ATAC-seq signal tracks show numerous accessible regions across the gene body. Regions whose enhancer activity is imaged in (C) are highlighted. B. Aggregate reads across the *fru* locus in *fru*^+^ versus *fru*^-^ male neurons plotted in 5kb windows. Pervasive opening is observed across much of the >100kb gene, with up to four-fold more reads in the *fru*^+^ condition. C. Peak landscapes and conservation across four genomic fragments in the Vienna Tiles collection, as indicated. Corresponding genomic locations are highlighted in A. D. Maximum intensity 2-photon stacks of enhancer activity of the four tiles in (C) within *fruitless* neurons in the male overlaid with the *fru*^P1^-LexA expression pattern (top) and as GFP signal alone (bottom). Each tile drives expression in distinct *fruitless* neurons. Remarkably, VT043701, which has higher ATAC signal in *fru*^-^ cells, drives expression in γ Kenyon cells, which we sorted into the *fru*^-^ population.

**Figure S3.**
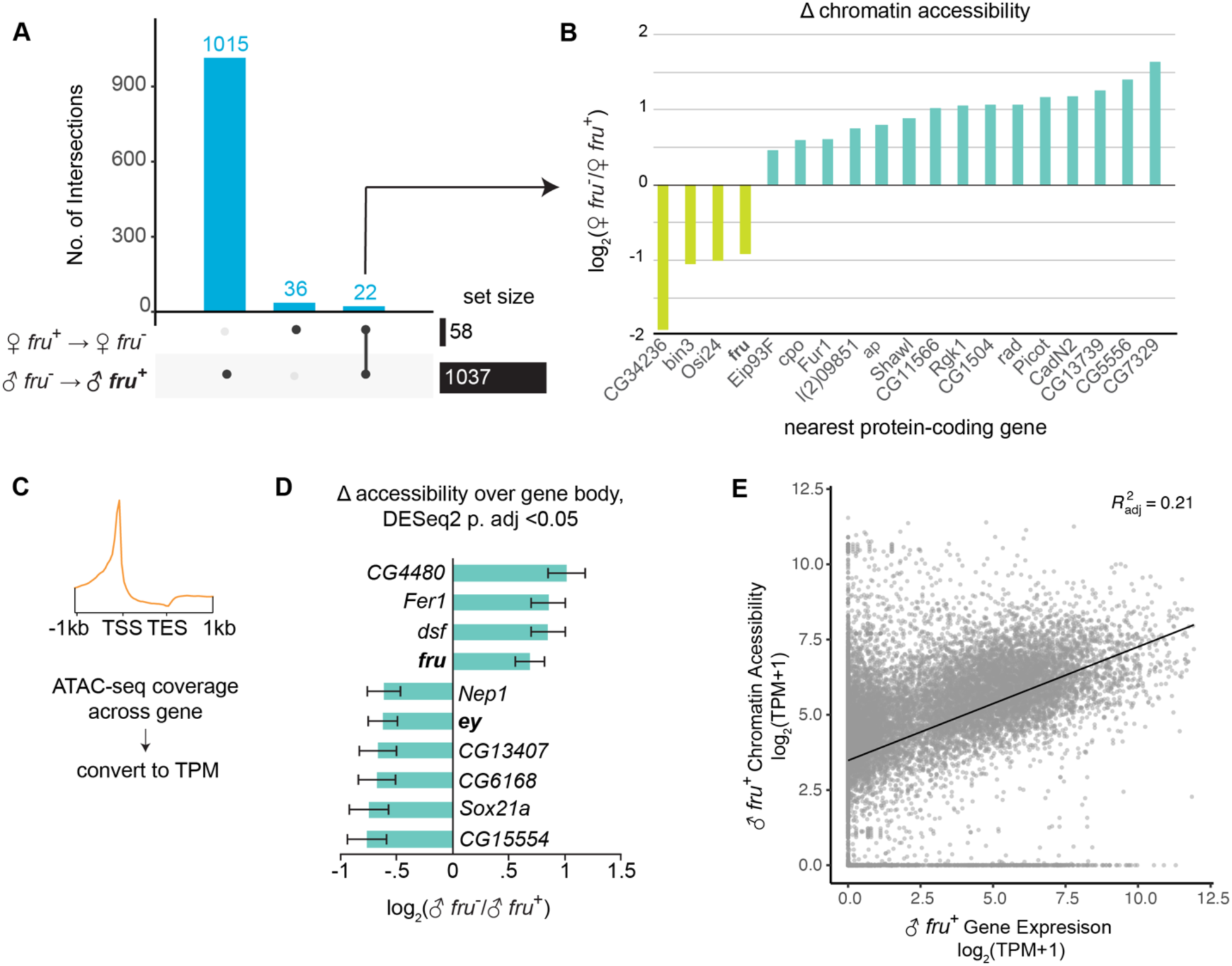
Chromatin accessibility changes in neurons based on transcriptional status of fruitless. A. UpSet plot showing the intersection of regions which are differentially accessible (DiffBind FDR <0.05) between *fru*^+^ and *fru*^-^ neurons in both sexes. B. Of 22 regions differentially accessible between *fru*^+^ and *fru*^-^ neurons, 19 correspond to protein-coding genes (labeled on the X axis). The log2(Fold Change) in accessibility of these genes relative in female neurons is plotted. C. Schematic of calculating chromatin accessibility across a gene locus. D. Barplot of log2(Fold Change) of genes with gene-scale differential coverage between male *fru*^+^ neurons and male *fru*^-^ neurons. *ey* is a Kenyon cell marker. E. Scatterplot of whole-gene chromatin accessibility versus gene expression level in the male *fru*^+^ dataset.

To ask whether this enhanced gene-wide coverage was unique to *fruitless*, we used featureCounts to count reads across a gene locus and used DESeq2 to measure changes in coverage between sample conditions using a cutoff of p.adj <0.05 (Figure S3C, D). *fruitless* was near the top of the list of genes with differential coverage in this analysis. We manually inspected peak landscapes of other high-ranked genes and found that most were short genes dominated by localized, robust peaks (like those found in *rox1* and *rox2*). We used this gene-scale ATAC coverage measurement to compare accessibility and RNA expression levels in the male *fru*^+^ condition and found mild correlation (r2=0.21, Figure S3E)

To ask if diverse regions of the *fru* locus act as active enhancers in different subpopulations of *fru*^*+*^ cells, we obtained reporter lines tiling upstream introns. These reporters consist of 1-3kb pieces of genomic DNA placed upstream of a minimal promoter and the coding sequence of the Gal4 transcription factor (47). We used a genetic intersection approach to identify *fru* neurons in which these fragments of the *fru* locus could act as enhancers (Figure 3C, D). Indeed, each region we examined drove expression in distinct *fruitless* subpopulations, and none of the regions we examined was able to drive gene expression across all *fruitless* neurons. We therefore conclude that the *fruitless* locus is densely packed with enhancer elements that each drive *fru* transcription in a subset of *fru*^+^ cells, i.e. that distinct gene regulatory mechanisms control *fru* expression across the diverse neurons that express it. This is consistent with analyses of the transcriptional control of neurotransmitter systems across ontogenetically diverse neuronal populations in *C. elegans* (48).

### Identification of candidate Fru^M^-regulated genomic elements

We sought to identify genomic regions whose accessibility robustly correlated with Fru^M^ status: Peaks present in male *fru*^*+*^ cells and absent from all other datasets represent genomic regions opened in the presence of Fru^M^, while peaks absent in male *fru*^*+*^ cells and present in all other datasets represent regions closed in the presence of Fru^M^. To identify these regions, we used DiffBind to call differentially accessible peaks between male and female samples, and between *fru*^*+*^ and *fru*^*-*^ samples (Figure 4A-C, Figure S4A-C). We observed 1037 differential peaks between male *fru*^+^ and male *fru*^-^, 861 between male *fru*^+^ and female *fru*^+^, and only 58 between the two female datasets (Figure 4A). The depletion of differential peaks between the two female datasets, both lacking Fru^M^, supports our hypothesis that the female *fru*^+^ cells, lacking Fru^M^, are a heterogeneous population whose constituents are no more similar to one another than those in the *fru*^-^ population. Further, we attributed a small number of chromatin accessibility changes which depend on sex or *fruitless* transcriptional status alone, suggesting that the large number of accessibility changes depend on the activity of Fru^M^ itself (Fig 4A).

**Figure 4.**
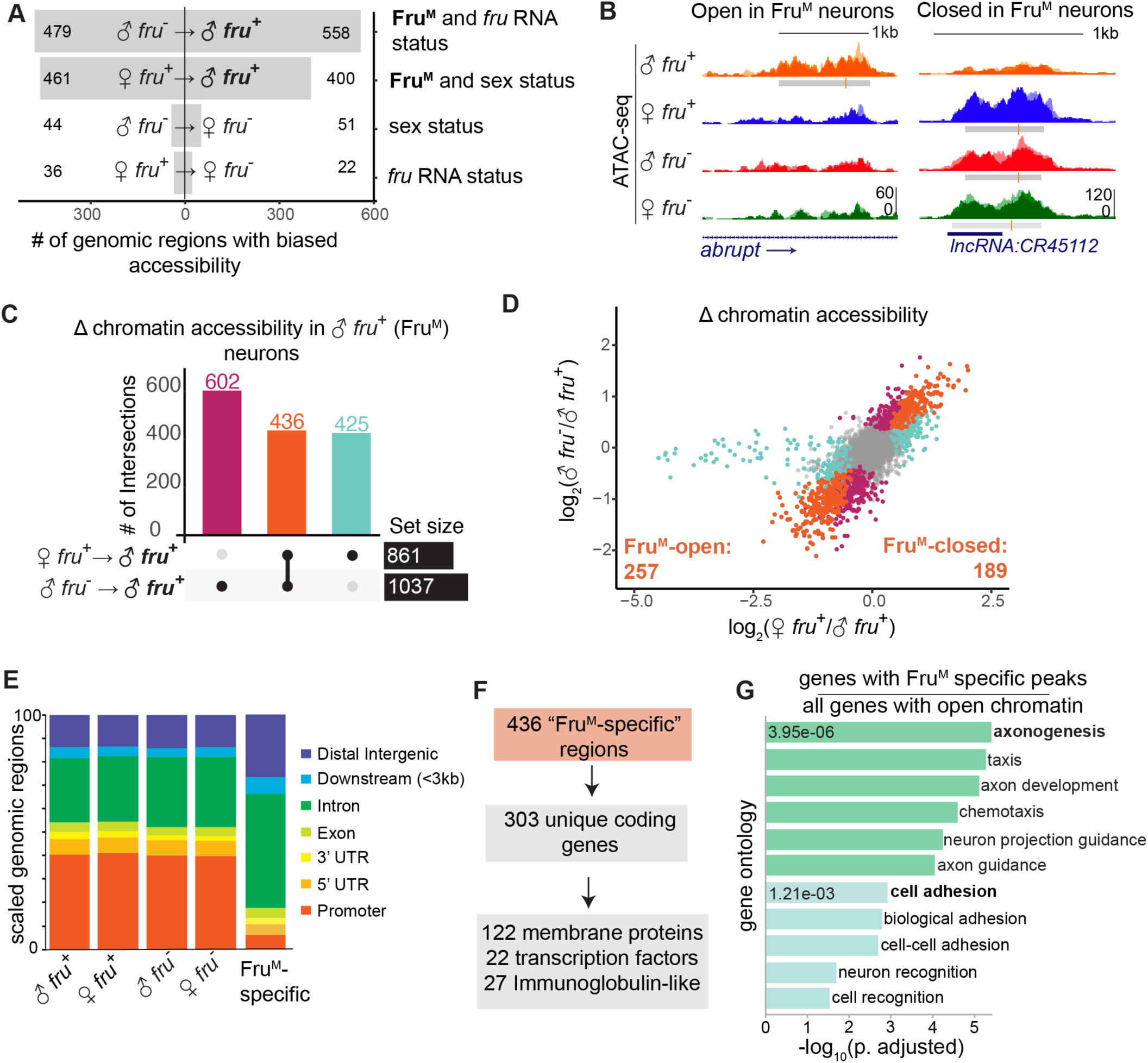
Chromatin changes downstream of Fru^M^ are near genes involved in neuronal projection and synaptic matching. A. Number of differentially accessible peaks between different sample comparisons at FDR<0.05. A “biased” region indicates a region which is relatively more open in a given comparison. Labels at right describe how compared conditions differ. B. Examples of peaks with increased (left) and decreased (right) accessibility specific to Fru^M^ neurons. C. UpSet plot showing intersection of DiffBind results at FDR <0.05 from male versus female *fru*^+^ neurons (861 sites) and male *fru*^+^ versus *fru*^-^ neurons (1037 sites). D. Distribution of fold change between the binary comparisons shown in (C). Each point is a peak. Points are colored according to their status in (C). All peaks that are differentially accessible in both comparisons vary in the same direction in both comparisons. Tail of aqua points at left represent dosage compensation signals from the X chromosome. E. Distribution of peak locations for all peaks versus Fru^M^-specific peaks. F. 436 Fru^M^-specific peaks are located near 303 unique genes, which are enriched for the gene categories noted. G. GO analysis using gProfiler of the 303 genes shown in (F). We include only terms with <1000 members. Genes with Fru^M^-specific annotated peaks were used as input gene lists, and all genes with peaks were used as background. Green terms relate to axonogenesis and teal to adhesion.

**Figure S4.**
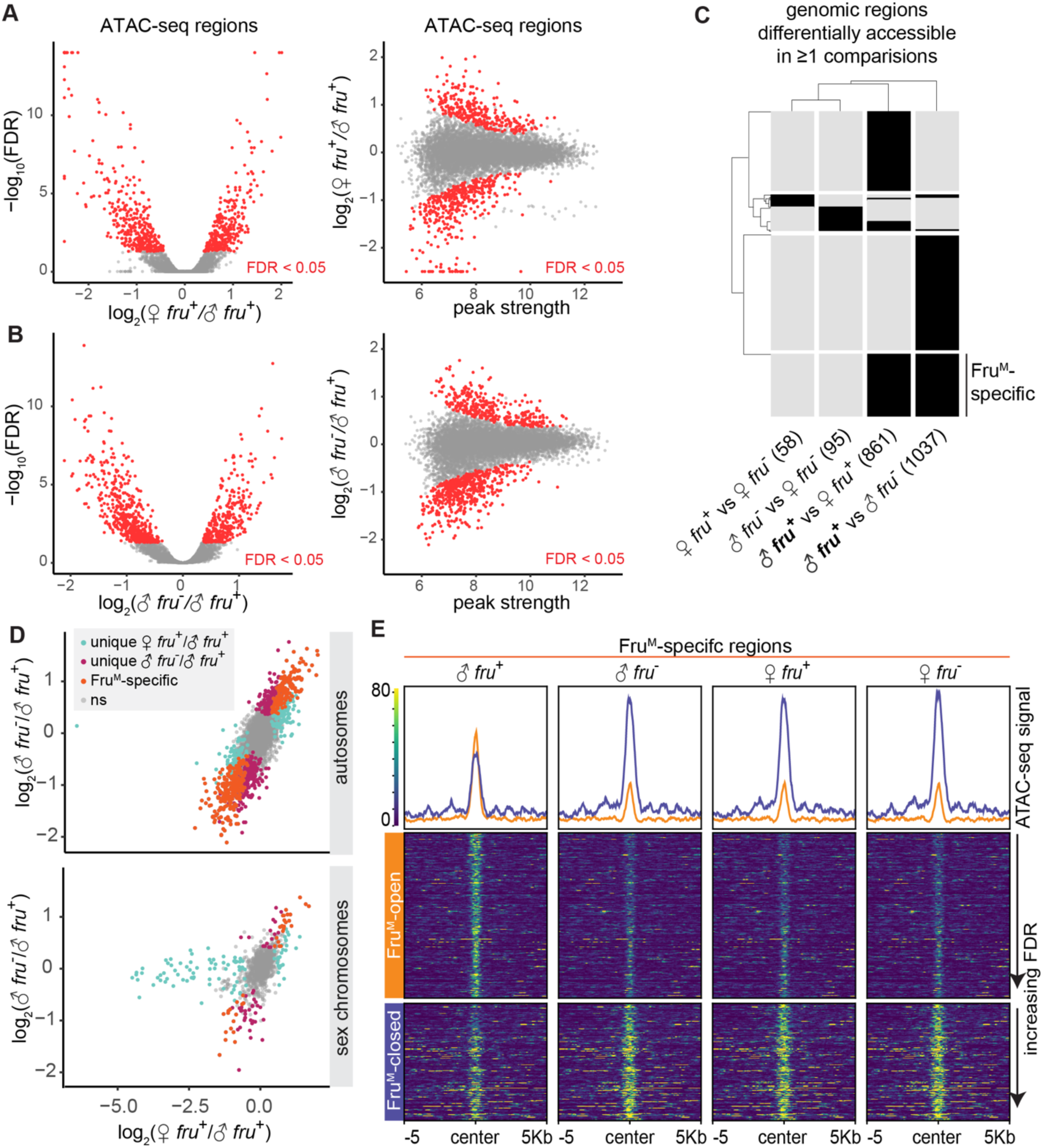
A. Volcano and MA plots of regions with differential accessibility (DiffBind < 0.05) between female and male *fru*^+^ neurons. Points at the top border in the volcano plot and along the bottom border in the MA plot have been thresholded such that they are visually comparable to plots in B. B. Volcano and MA plots of regions with differential accessibility (DiffBind < 0.05) between male *fru*^-^ and *fru*^+^ neurons. C. Binary heatmap of regions called differentially accessible in 1 or more comparisons. D. Scatterplot of log2 fold changes in accessibility compared to male *fru*^+^ neurons. Figure shows analysis in Figure 4D with autosomal regions (chromosomes 2, 3, and 4) separated from sex chromosomal regions (X and Y). E. Signal heatmaps of Fru^M^-specific regions, separated by regions which are selectively open or closed in male *fru*^+^ neurons.

To identify changes in the activity of gene regulatory elements downstream of Fru^M^, we took the intersection of (1) the 1037 peaks specific to male *fru*^+^ versus male *fru*^-^ and (2) the 861 peaks specific to male *fru*^+^ versus female *fru*^+^ (Figure 4C, D). This resulted in 436 high-confidence peaks (FDR<0.05 in both comparisons) genome-wide. Comparing between the two male samples allows us to filter out sex-specific, Fru^M^-independent elements, while comparing between male and female *fru*^+^ cells allows us to compare populations of cells with roughly matched identities and thus filter out peaks associated with cell type distribution. In line with our biological expectations, all peaks that satisfied both conditions were biased in the same direction in both comparisons i.e. present in the male *fru*^+^ dataset and absent in the other three (257 peaks), or absent in the male *fru*^+^ dataset and present in the other three (189 peaks) (Figure 4D, Figure S4D). Most differential peaks were intronic or intergenic, while promoters were sharply underrepresented among differential peaks (Figure 4E). We consider these 436 genomic elements to be candidate enhancers or repressors whose activity is regulated directly or indirectly by Fru^M^.

The fly genome is compact relative to common model vertebrates, and regulatory elements are often found in or near the genes they regulate (47). We therefore hypothesize that our 436 candidate elements regulate the genes closest to them. Our Fru^M^-dependent peaks were in or near 303 unique genes, which were particularly enriched for membrane proteins (122 genes), transcription factors (22) and immunoglobulin superfamily (IgSF) members (27) (Figure 4F). GO terms for cell adhesion and axonogenesis were also highly enriched compared to all the genes containing peaks across our dataset (Figure 4G). Of course, these are attractive candidate factors for determination of neuronal identity or connectivity, and IgSF proteins have been implicated previously in the *fru* circuit (49).

We manually inspected a subset of the 436 candidate regulatory elements; they were clearly differentially accessible by both qualitative inspection and statistical thresholds, were reproducible between replicates, and were found in genes plausibly involved in the biological processes under study. However, these differential peaks were much smaller than the peaks we observed in promoters, which were of common height across samples (Figure 4E, and see *fruitless* locus in Figure 1D). Because “accessibility” as measured by ATAC is essentially quantized—each region of the chromosome is tagged 0, 1, or 2 times per cell—we reasoned that peak height provides a rough measure of the proportion of cells in the analyzed population in which a locus is open, or unbound by nucleosomes. If this is the case, we would predict that each regulatory element we identified is being used by only a small portion of the *fru*^+^ cell population, suggesting that Fru^M^ might induce different gene regulatory programs in different ontogenetic classes of *fru*^+^ neurons.

### Genomic elements specifically accessible in male *fruitless* neurons act as enhancers in subsets of male *fruitless* cells

To test the ability of candidate gene regulatory elements to drive gene expression, we identified reporter Gal4 alleles matched to their genomic loci, as in the *fru* locus shown in Figure 3C-E (47,50). Such reporter alleles are available for many of our 436 Fru^M^-dependent elements, and we selected the top seven ATAC-seq peaks specifically open in Fru^M^ cells for which reporter alleles were available (Figure 5A, D, G). In order to visualize neurons with recent transcription from the *fruP1* promoter, we constructed animals in which fruP1-LexA (51) drove expression of a red fluorophore, and reporter Gal4 constructs drove GFP expression. These animals allowed us to examine overlap between the *fruitless* population and neurons in which the candidate genomic region was capable of acting as a transcriptional enhancer (Figure 5B, C, E, F). To simplify observation of enhancer activity within the *fruitless* population, we also used an intersectional genetic approach to identify neurons positive for enhancer activity and current or past expression from *fruP1* (Figure S5A, B). Enhancer activity of all seven fragments was tested with both genetic strategies, with consistent results.

**Figure 5.**
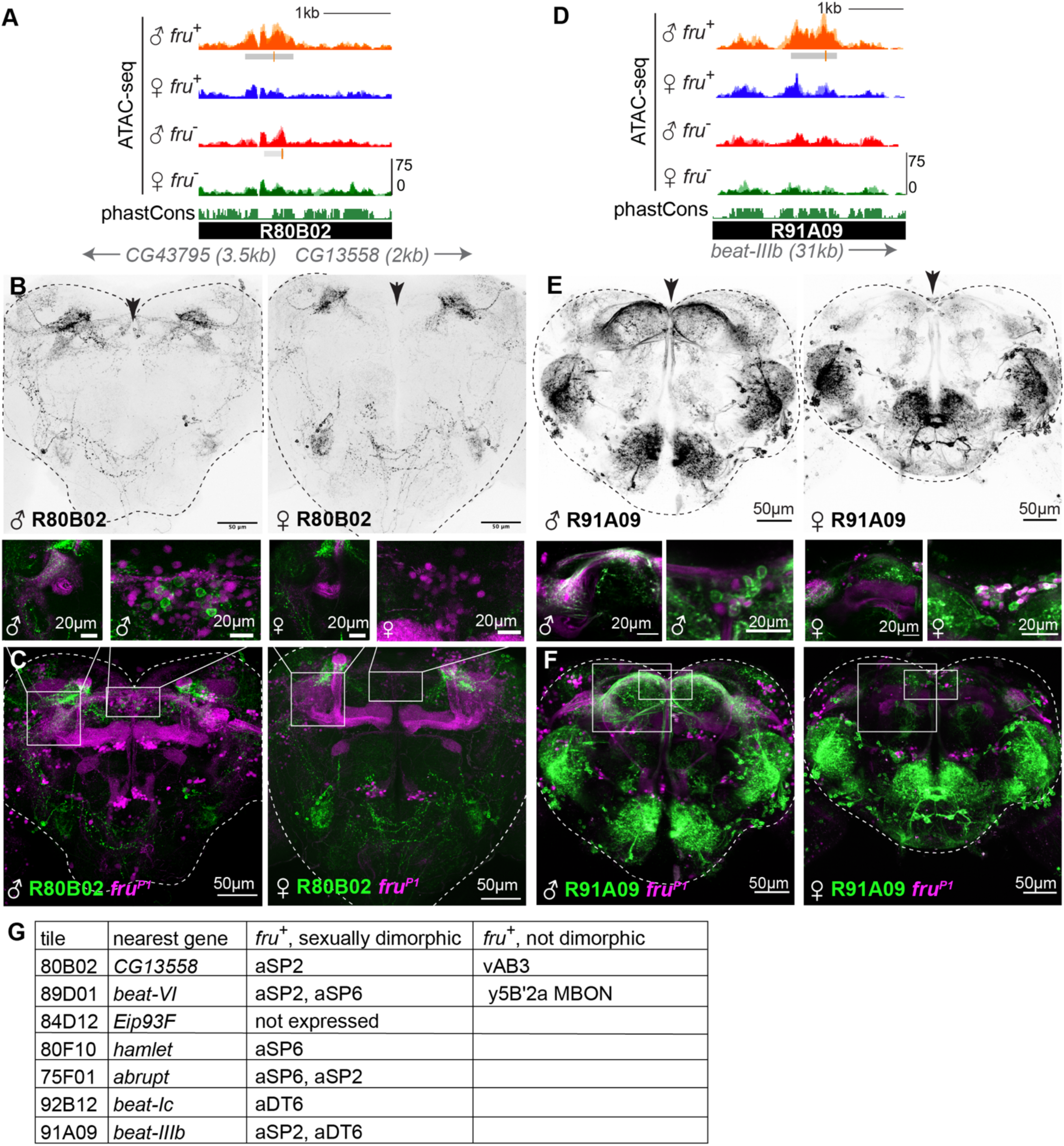
Regulatory elements specifically accessible in FruM neurons have sexually dimorphic and cell-type-specific activity *in-vivo*. A. UCSC genome browser screenshot of an intergenic region with accessibility specific to Fru^M^ neurons. The 80B02 fragment encompasses this peak. B. Confocal maximum intensity projections of reporter element 80B02 driving 10XUAS-IVS-myr::GFP in male (left) and female (right). 80B02 activity is sexually dimorphic in cells with somata located at the dorsal midline (arrowhead), and common to the sexes in dorsolateral cells that innervate the central complex. Representative of 3-4 images per sex. Imaging conditions were matched between sexes. C. Overlap of 80B02 signal with *fru*^*P1*^-LexA-driven myr::TdTomato expression in the brains shown in (B). Insets show double-positive somata at the male dorsal midline and fibers in the lateral protocerebral complex. D. UCSC genome browser screenshot of a second intergenic region with accessibility specific to Fru^M^ neurons. The 91A09 fragment encompasses this peak. E. Maximum intensity projection of two-photon stack of 91A09-Gal4 driving 10XUAS-IVS-myr::GFP in male and female. 91A09 expression is shared between the sexes in lateral and subesophageal regions, but male-specific dorsally (arrowhead). Representative of 2-3 images per sex. Imaging conditions were matched between sexes. F. Overlap of 91A09 signal with *fru*^*P1*^-LexA-driven myr::TdTomato expression in the brains shown in (B). Insets show double-positive somata at the male dorsal midline and fibers in the lateral protocerebral complex. G. Summary of *fru*^+^ cell types in which analyzed genomic fragments drive reporter expression.

**Figure S5.**
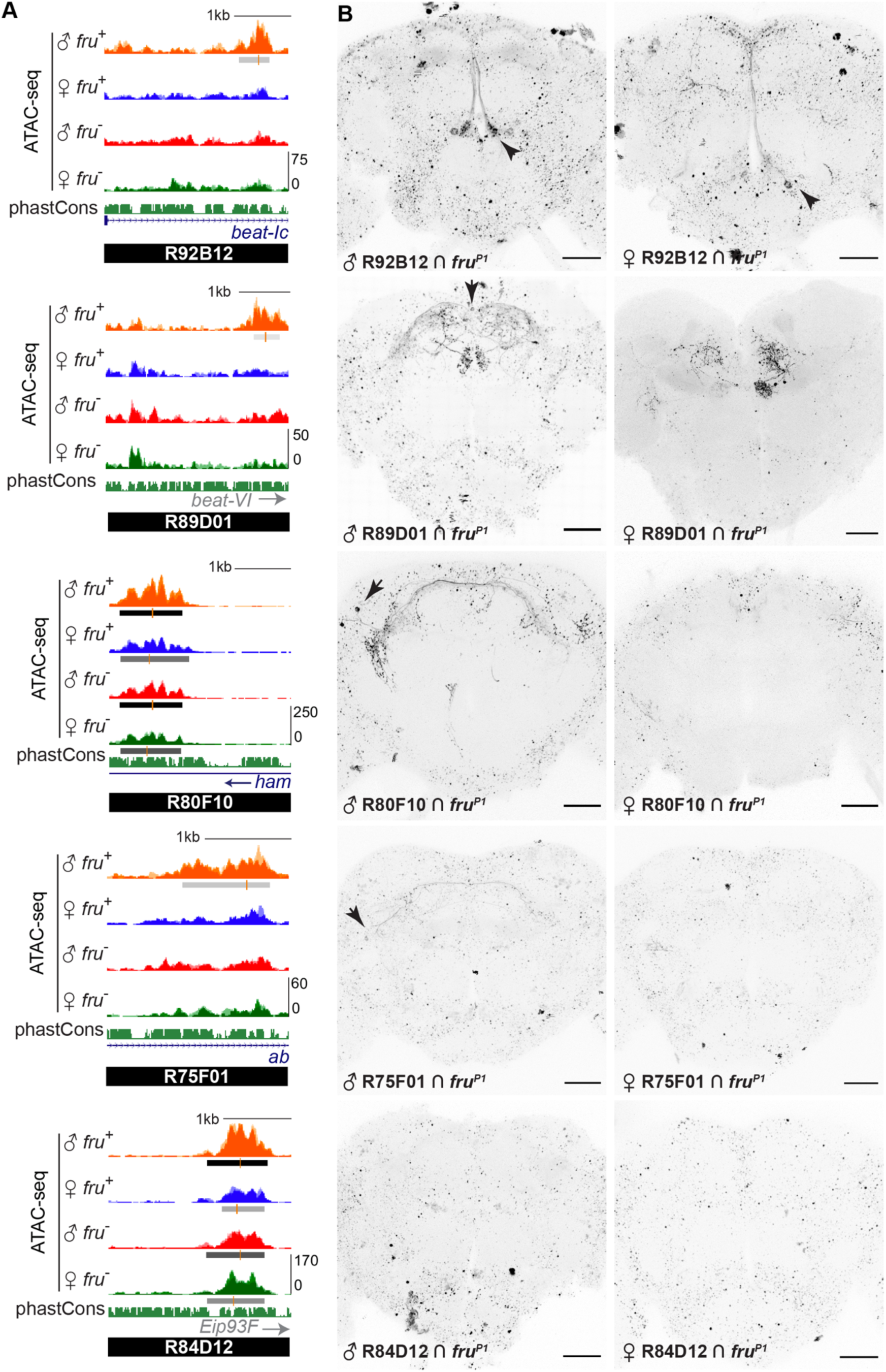
Sexually dimporphic enhancer usage in distinct Fru^M^ neurons. A. UCSC genome browser screenshots of genomic regions covered by each reporter. B. 2-photon stack of adult male (B) and female brains. Arrows point to cells with sex-specific expression (89D01, 80F10, 75F01) or dimorphic labeling intensity (92B12). 84D12 labeling is mutually exclusive with *fru* expression. Brain-wide speckle signal is autofluorescence.

In six of seven cases, we observed male-specific reporter expression in one or a few anatomic classes of *fruitless* neurons, validating these differential peaks as bona fide enhancers with sex-specific activity and confirming that our ATAC-seq peaks demarcated accessible chromatin from subpopulations of *fru*^+^ neurons (Figure 5 B, C, E, F, G, Figure S5B). Reporter expression driven by the seventh construct was mutually exclusive with fru-LexA expression (84D12 tile, Figure S5B). In addition to sex-specific labeling of *fru*^*+*^ neurons, each allele also drove non-dimorphic reporter expression in some *fru*^*-*^ neurons (e.g. as seen in Figure 5C, F). Because reporter tiles were much larger than our peaks (2-3 kb versus 500bp), they could comprise multiple enhancer elements, only one of which is Fru^M^-dependent. Alternatively, minimal regulatory elements could be pleiotropic, such that they can be bound by alternative trans-acting factors in the absence of Fru^M^. Future enhancer-bashing experiments will be required to discriminate between these possibilities. We also note that three classes of *fruitless* neurons, aSP2, aSP6, and aDT6, were repeatedly labeled by these six fragments. These *fruitless* classes are particularly numerous, and we assume their over-representation reflects the fact that we analyzed reporters matched to the strongest differential peaks. Reporters never labeled all the aDT6, aSP2, or aSP6 neurons, suggesting that these anatomic groups contain multiple transcriptional subtypes.

Finally, while most reporter constructs drove sex-biased expression in subpopulations of male *fruitless* neurons, none of the reporters drove expression across the whole *fruitless* population. Together with the scale of the differential peaks we observe, these data lead us to conclude that Fru^M^ has different direct and/or indirect genetic targets in different ontogenetic/anatomic subpopulations of *fruitless* neurons. Moreover, individual Fru^M^-regulated enhancers are activated downstream of Fru^M^ only in specific subpopulations of *fruitless* neurons.

### Transcriptional effectors of Fru^M^ are neither universal across *fru*^+^ cells nor dedicated to *fru*^+^ cells

Our ATAC-seq and enhancer activity assays suggest that Fru^M^ genomic targets vary across distinct anatomic populations of *fruitless* neurons. To ask whether there are any genes broadly regulated by Fru^M^ status across the *fruitless* population, we used DESeq2 on our RNA-seq data to call differentially expressed genes between male and female *fru*^*+*^ neurons, between male *fru*^*+*^ and *fru*^*-*^ neurons, and between female *fru*^*+*^ and *fru*^*-*^ neurons (Figure 6A, S6A, B) (52). As shown in Figure 1, *fruitless* transcript quantity and splice isoform tracked the cell type and sex of the library. Aside from *fruitless*, we observed <300 genes differentially expressed between male *fru*^*+*^ and male *fru*^*-*^ cells, and most of these were also differentially expressed, in the same direction, between female *fru*^+^ and female *fru*^-^ cells (Figure 6A, S6A, B). We interpret these to be signatures of the particular populations of cells we analyzed, and that these differences in expression at the population level are independent of Fru^M^. For example, *eyeless* is a marker of Kenyon cells, which we sorted into the *fru*^*-*^ population in both sexes; *eyeless* transcripts are enriched in both *fru*^*-*^ datasets, as expected. Many other genes common to *fru*^+^ or *fru*^-^ datasets were involved in neurotransmitter or neuropeptide production or reception suggesting the potential for distinct distributions of transmitter usage between sexually dimorphic (*fru*^+^) versus sex-shared (*fru*^-^) cells (Figure S6C).

**Figure 6.**
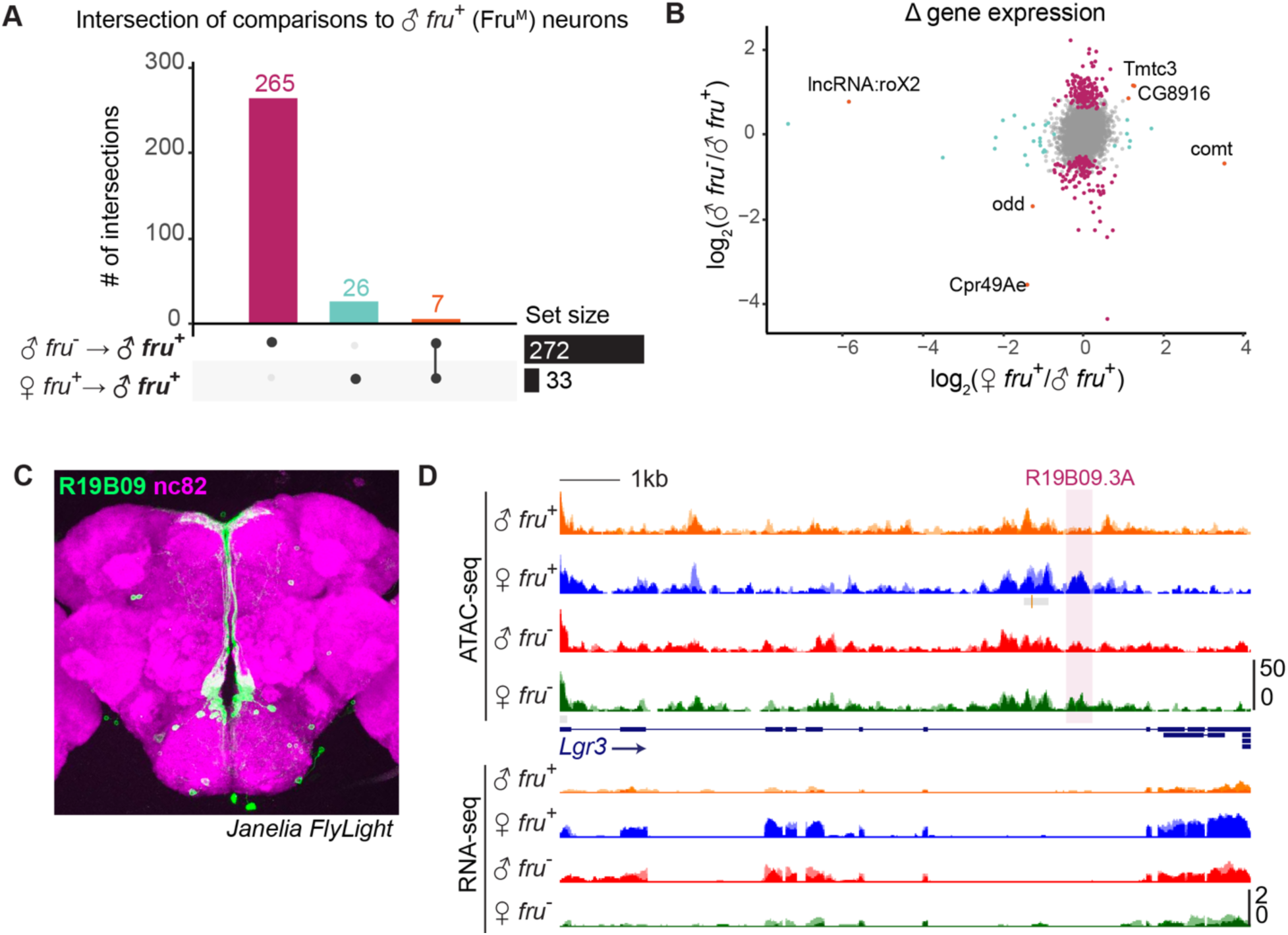
Fru^M^-regulated genes are neither distinct to nor universal across male *fruitless* neurons. A. UpSet plot showing intersection of differential gene expression (DESeq2 results at FDR <0.05) from male versus female *fru*^+^ neurons (33 genes) and male *fru*^+^ versus *fru*^-^ neurons (272 genes). Only 7 genes are differential across both comparisons. B. Distribution of fold change between the binary comparisons shown in (A). Points are colored according to their status in (A). Only 5 genes have differential expression specific to Fru^M^ cells (i.e. fold changes in the same direction in both comparisons). C. Adult female brain expression pattern of R19B09 (green) driving expression in aDT6/median bundle neurons with nc82 counterstain (magenta). image from Janelia FlyLight database. D. UCSC genome browser screenshot of RNA-seq and ATAC-seq signal at *Lgr3*, a gene that is expressed in female but not male median bundle neurons (25). Highlighted is R19B09.3A, which Meissner et al. found to be a minimal regulatory element bound by Fru^MB^.

**Figure S6.**
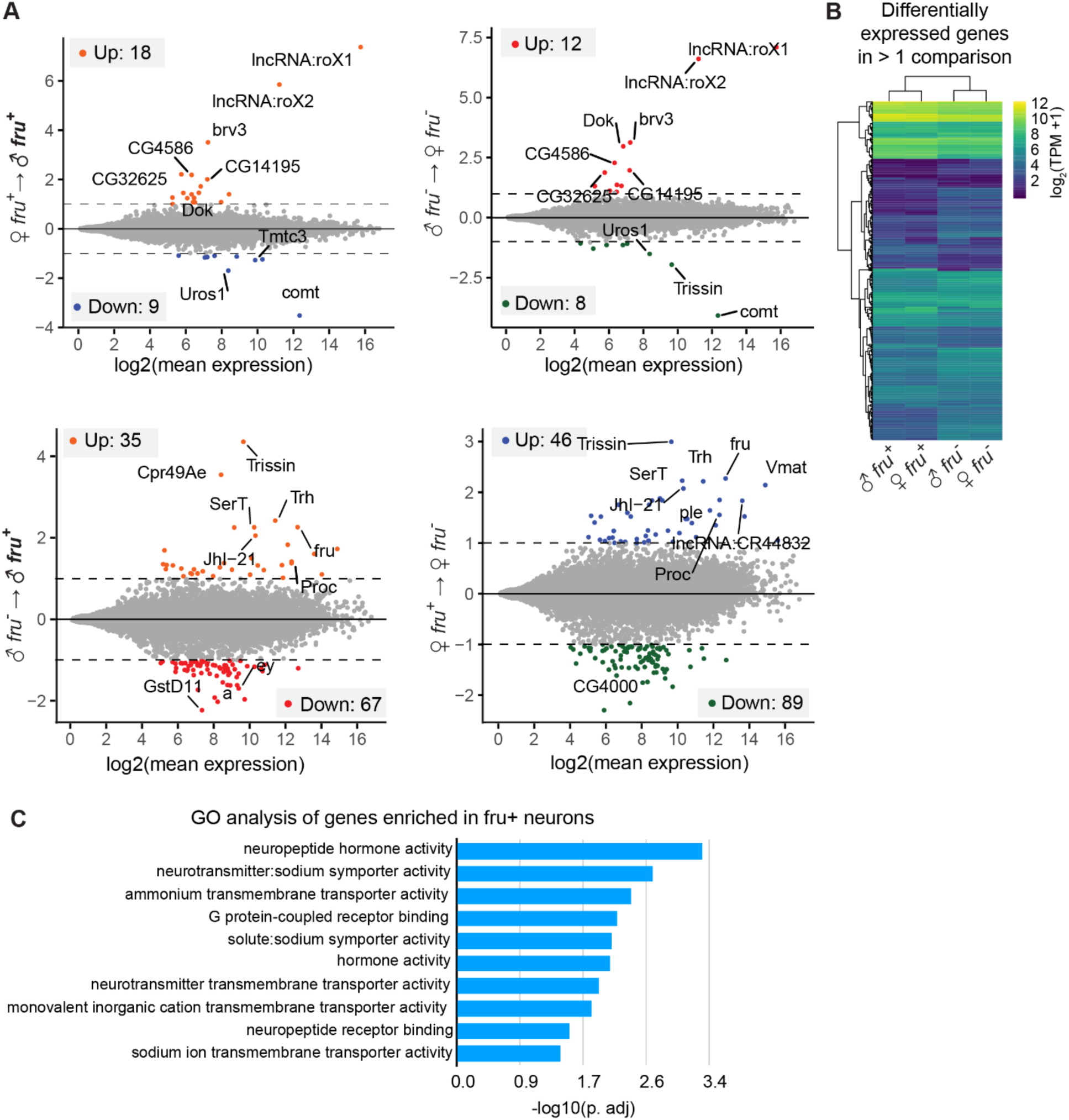
A. MA plots of differential RNA expression analysis between four datasets. B. Clustered heatmap of TPM values of genes with differential expression in one or more comparisons. Differential expression is dominated by *fru* status. C. Gene ontology analysis of genes enriched in both *fru*^+^ datasets (male and female) over *fru*^-^ datasets. Enrichment is over a custom background of genes with expression > 10 TPM across the four datasets.

We observed only 33 statistically significant differences (DESeq p. adj <0.05) in gene expression between male and female *fru*^*+*^ neurons, most of which were also differed between male versus female *fru*^*-*^ neurons (Figure S6A). We interpret these as being sex-specific but Fru^M^ independent.

If Fru^M^ alters transcription of distinct genes across the dozens of classes of *fruitless* neurons, we would expect these differences to average out when the whole pool of *fruitless* neurons is analyzed *en masse*. In contrast, if Fru^M^ regulated the same effectors across all *fruitless* classes, we would expect to observe a strong transcriptional signature specific to male *fru*^+^ cells, containing Fru^M^. We identified only seven transcripts that are uniquely active or inactive in cells containing Fru^M^ (Figure 6A, B). The implication of this analysis is that aside from *fruitless* itself, there are unlikely to be strong differences in gene expression that are dedicated to or shared across the *fru*^*+*^ cell population in the adult. These findings are consistent with our results that Fru^M^ regulates different genomic elements in different populations of *fru*^*+*^ cells and with prior research suggesting that the same neuronal specificity factors are used throughout the brain in different combinations (53–55).

Fru^MB^ was shown previously to bind to a minimal regulatory element in *Lgr3* and to repress *Lgr3* expression in a subset of male Fru^M^ median bundle/aDT6 neurons (Figure 6C) (25). Remarkably, we observe increased transcription of *lgr3* in pooled female *fru*^+^ cells, which we presume emanates from the aDT6 subpopulation; aDT6 cells comprise 5-10% of our *fru*^+^ populations (Figure 6D). Fru^MB^ could decrease *Lgr3* expression in the male by decommissioning an *Lgr3* enhancer or commissioning an *Lgr3* repressor. To discriminate between these, we visualized this minimal region in our ATAC-seq data and found it to be accessible in female *fru*^+^ cells and not in male *fru*^+^ cells (Figure 6D). This suggests that the regulatory element in *Lgr3* identified by Meissner et al. is an enhancer that is decommissioned by Fru^M^.

### Enrichment of Fru^M^ binding sites in peaks specifically closed in male *fru*^+^ neurons

To ask whether Fru^M^ acts to decommission regulatory elements more generally, we first needed to separate direct targets (i.e. those bound by Fru^M^), from indirect targets (i.e. those regulated by direct Fru^M^ targets). To explore this using our ATAC data, we searched for Fru motifs across the 436 Fru^M^-specific peaks. Fru^M^ has three DNA-binding domains, termed Fru^A^, Fru^B^, and Fru^C^. The three isoforms are expressed in largely overlapping neuronal populations, with Fru^B^ and Fru^C^ expressed more broadly than Fru^A^ (30,56). The Fru^B^ and Fru^C^ isoforms of Fru^M^ are each independently required in the male for courtship behavior, while loss of Fru^A^ has little effect on courtship (30,56). Loss of Fru^C^ causes feminization of neuronal anatomy, while loss of Fru^A^ and Fru^B^ have little anatomic effect (30). The three isoforms have been shown by SELEX to bind distinct DNA motifs (Figure 7A, Figure S7A) (57).

**Figure 7.**
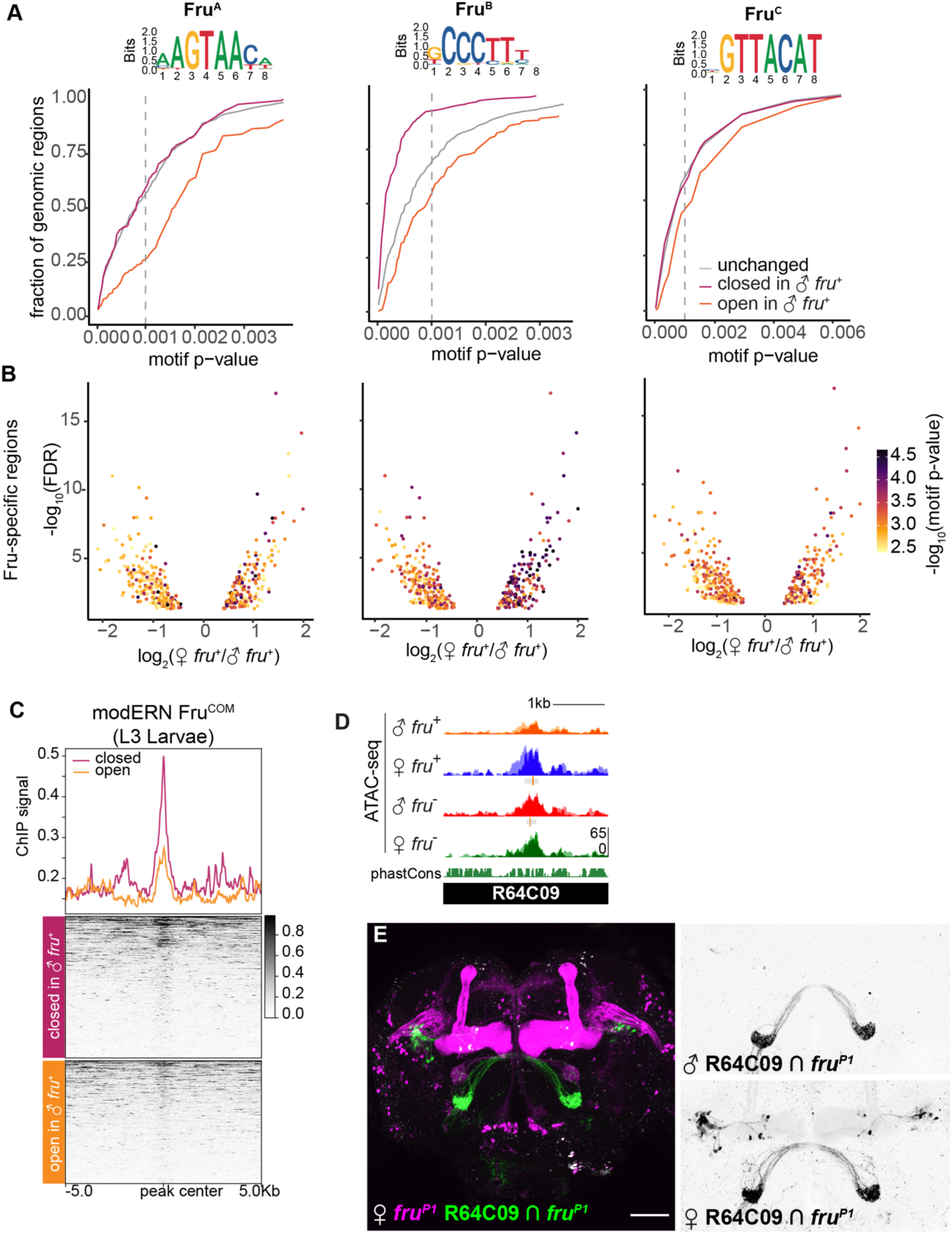
Strong FruM motifs are enriched among peaks closed in the presence of Fru^M^. A. SELEX motif for Fru^B^ (57) and cumulative frequency distribution of Fru^B^ motif strengths identified using FIMO across peaks open (orange), closed (magenta), and unchanged (grey) in the presence of Fru^M^. All three motifs are depleted from regions specifically open in Fru^M^ neurons, and Fru^B^ motifs are enriched among peaks specifically closed in the presence of Fru^M^. The p-value threshold for a well-matched motif (p <0.001) is marked as a dashed line. B. Volcano plots showing the strength of the best match to each motif across our 436 Fru^M^-specific peaks. Regions opened in the presence of Fru^M^ have negative values. C. Fru^COM^ binding profiles across our Fru^M^-closed and Fru^M^-opened peaks. Fru^COM^ data is from whole L3 larvae (60). Fru^COM^ signal is enriched at Fru^M^-specific peaks, especially those closed in the presence of Fru^M^. D. UCSC genome browser screenshot of ATAC-seq signal across a FruM-closed region covered by the enhancer reporter element R64C09. E. Genetic intersection of R64C09 enhancer activity with *fruitless* expression. R64C09 labels a distinct neuron population in female that is not labeled in male *fruitless* neurons, consistent with it acting as an enhancer that is decommissioned in the presence of Fru^M^. 64C09 also drives sex-shared expression in *fru*^+^ olfactory sensory neurons.

**Figure S7.**
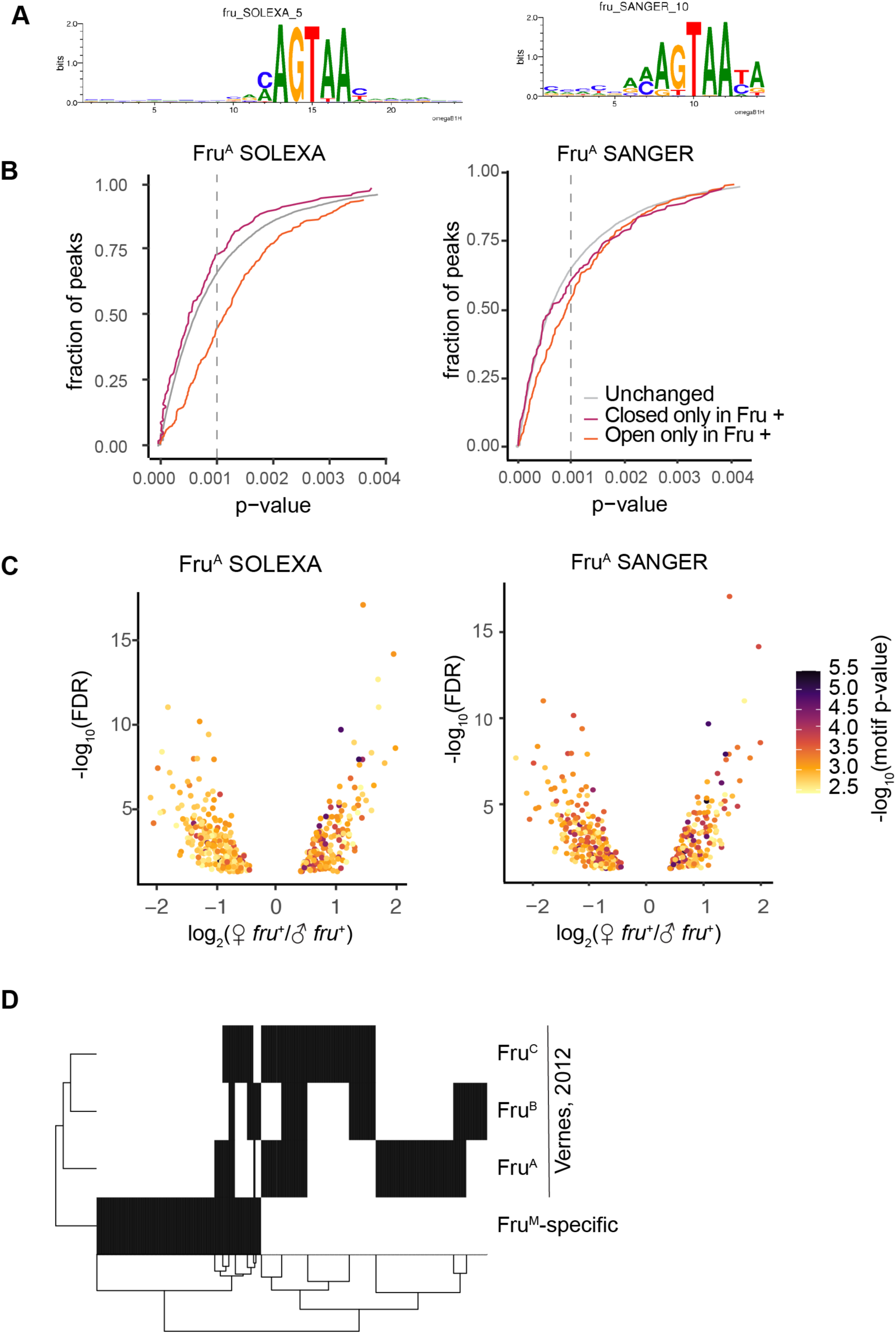
Strong Fru^M^ motifs are enriched among peaks closed in the presence of Fru^M^. A. SELEX and SANGER Fru^A^ motifs. B. Cumulative frequency plots for SELEX and SANGER Fru (Fru^A^) motifs (FlyFactorSurvey) across Fru^M^-open, Fru^M^-closed and unchanged regions. C. Volcano plots showing motif strengths across Fru^M^-specific regions. D. Binary heat map of overlap of Fru^M^-specific regions with previously identified Fru^M^ targets in S2 cells (61).

We used FIMO to search our peak datasets for Fru^A^,^B^, and ^C^ motifs. In each peak, we quantified the strongest match to each of these motifs (58). We were surprised to find that motifs for all three DNA binding domains were more common among peaks specifically closed in the presence of Fru^M^ than in those specifically open in the presence of Fru^M^ (Figure 7A, B, Figure S7A-C). This pattern was particularly apparent for Fru^B^ motifs, which were strongly enriched in peaks closed in the presence of Fru^M^ (Figure 7A, B). Together, these results suggest that, in the adult, Fru^M^ decommissions (or closes) the regulatory elements to which it directly binds. Moreover, these results suggest that the 189 regions inaccessible in male *fru*^*+*^ cells are those most likely to be direct Fru^M^ targets, while regions specifically opened in male *fru*^*+*^ cells are likely to be downstream effectors of primary targets. Analysis of additional Fru^A^ motifs shown in Figure S7A-C.

Fru^M^ and Fru^COM^ share most of their protein coding regions and their DNA binding domains, and Fru^COM^ can rescue loss of Fru^M^ in behavioral experiments (59). We therefore asked whether our putative Fru^M^-regulated sites overlapped with Fru^COM^-bound sites identified by ChIP from larvae through the modERN project (ENCODE ENCGM860XOW) (60). We re-analyzed raw data from the modERN dataset and compared ChIP enrichments across our peaks closed in the presence of Fru^M^ versus open in the presence of Fru^M^. We found strong enrichment of Fru^COM^ binding centered over peaks closed in the presence of Fru^M^, and weaker Fru^COM^ binding enrichment at peaks opened in the presence of Fru^M^ (Figure 7C). This analysis suggests that despite distinct cellular contexts, Fru^M^ and Fru^COM^ may have targets in common, and supports our hypothesis that Fru decommissions regulatory elements to which it directly binds.

To test whether additional regions closed in Fru^M^ cells function as enhancers in female *fruitless* neurons, we selected one such region, an intergenic peak located near *cry, vib*, and *CG31475* (Figure 7D). We used our intersectional genetic strategy to analyze expression of 64C09, a Janelia fragment encompassing this peak (Figure 7E). Remarkably, we found that this element drove robust, female-specific reporter expression in a subset of *fru* neurons. Together, these results suggest that Fru^M^ acts cell-type-specifically to decommission enhancer elements in male *fruitless* neurons.

## Discussion

Here, we have analyzed the landscape of gene regulatory elements upstream and downstream of the *fruitless* transcription factor. Together, our results suggest that Fru^M^ is a single node of commonality across cells that are otherwise transcriptionally diverse. The mechanisms upstream of *fru* transcription are likely to be distinct across different cells; once Fru^M^ is translated in males, it does not unify gene expression programs across the disparate cells that express it, but rather executes distinct programs in each type of *fru*^+^ cell. Fru^M^ therefore serves as an evolutionary and developmental handle on neurons that need to be made sexually dimorphic, and likely intersects differently with the gene regulatory identities of the individual neuron types in which it is expressed to alter them in different ways.

### Fru^M^ as a decommissioner of regulatory elements

Despite decades of work on the cellular, circuit, and behavioral functions of Fru^M^, we know very little about what Fru^M^ does as a transcription factor to masculinize neurons. Our findings suggest that Fru^M^ directs genome-wide changes in chromatin accessibility and decommissions its direct targets. This mechanism is strikingly consistent with previous work: The two genes previously identified as direct Fru^M^ targets, *robo1* and *Lgr3*, are both downregulated in the male (25,26). Moreover, Fru^M^ can recruit the cofactor Bonus, followed by the chromatin modifiers HDAC1 or HP1 (Ito et al., 2012); HDAC1 and HP1 both act to convert euchromatin to heterochromatin.

This decommissioning activity raises an important question of whether Fru^M^ remains bound to such loci. Currently, few transcription factors show ability to bind nucleosomes (“pioneer activity”). And while currently no high-resolution binding profiles exist for Fru^M^, ChIP enrichment of Fru^COM^ in whole third instar larvae correlates with accessible chromatin in third instar larval brains. This suggests Fru proteins bind nucleosome-displaced and not nucleosome-occupied regions. FruM may then use a “hit and run” mechanism – binding a locus, recruiting chromatin modifiers, and leaving when the region is no longer accessible. If so, ChIP-based methods may miss direct targets of Fru^M^ which can nevertheless be detected by looking at resultant chromatin changes with ATAC-seq.

Beyond a putative molecular mechanism for the action of Fru^M^ on gene expression, this finding suggests that neuronal masculiization in the insect proceeds through cell-type-specific dismantling of female or sex-shared gene expression programs. Maleness is typically conceived as an addition to female programming, as exemplified by the striking male-specific courtship routines performed in birds and insects. However, Fru^M^ is not required for courtship actions per se, but rather for their regulation (11,62). Moreover, *fru*-expressing neurons in the female control egg-laying, and ectopic Fru^M^ expression in female flies dismantles egg-laying behavior (15,63). Together, these findings are consistent with masculinization as a process of loss of female programming.

### Fru^M^ masculinizes transcription differently in different classes of *fruitless* neurons

Fruitless masculinizes dozens of classes of neurons in the central brain alone, and does so by altering cell genesis, apoptosis, arbor anatomy, connectivity, and neurophysiology (2,3,6,20,21,31,32). Consistent with the variable nature of these masculinizing mechanisms, we find that Fru^M^ status alters the activity of distinct gene regulatory elements across distinct classes of *fruitless* neurons. Indeed, we find that each relevant genomic region is likely Fru^M^- regulated in just one or a few *fruitless* populations.

The specificity of Fru^M^ action across distinct *fruitless* neurons could result from distinct availability of cofactors with different DNA-binding domains, or from differences in the pre-existing chromatin landscape that Fru^M^ encounters in each cell type. Fruitless is one of 40 members of the BTB/POZ family of transcription factors in *Drosophila*. The BTB/POZ domain is a dimerization domain, and family members have been shown to both homo- and hetero-dimerize. The presence of different Fru^M^ heterodimers across neuron types that bind different composite motifs is thus a feasible mechanism. Another interesting possibility is that the distinct factors that activate *fruitless* transcription in different populations of *fruitless* neurons could themselves cooperate with Fru^M^ in regulating cell-type-specific effectors.

### Comparison with previous genome-wide datasets

Two previous studies attempted to identify Fru^M^-bound sites genome-wide through ectopic expression of tagged isoforms that enzymatically label DNA. Each of these experiments is conceptually difficult to interpret because Fru^M^ is studied outside of the typical cellular context and transcriptional millieu. A DamID study (56) analyzed central nervous systems in which Fru^M^-Dam expression is driven by leak from an uninduced UAS-driven construct, allowing expression in any cell at any time of development; a BirA/BLRP study (61) induced Fru^M^ expression in the S2 cell line, which is not neural (64). Reported binding is at the level of whole genes in the DamID study, and peaks are of median length 10kb in the BirA study. The low resolution, likely a result of the spatial resolution of enzymatic tagging and/or the accretion of covalent DNA labeling over long time scale labeling, preclude a crisp comparison with our nucleosome-scale peak landscapes. Gene-level intersection of putative Fru binding from the BirA/BLRP study (61) versus genes with differential peaks in our analysis is shown in Figure S7D.

We argue here that Fru^M^ alters expression of distinct genes across *fru* subpopulations, and that these transcriptional differences are masked when the whole *fru* populations are analyzed by RNA-seq. However, a previous study identified 772 genes enriched in male *fru*+ neurons in adult by TRAP (65). We were somewhat surprised to find that we did not re-discover these genes, especially as our analysis is more sensitive than TRAP (i.e. we observe 4.8-fold enrichment of *fru* versus 1.5-fold enrichment by TRAP). One possibility is that differences in TRAP were dominated by *fru*^+^ Kenyon cells, which we filtered out in our analysis (Figure 1A). The TRAP approach also relied on comparing polysome-bound transcripts from *fru*^+^ neurons with input from the whole head, which would be expected to enrich neural transcripts generally.

### Action of Fru^M^ across the life cycle

*fruitless* expression begins in late larvae, peaks in mid-pupal stages, and continues robustly in the adult; new populations of cells turn on *fruitless* expression across the life cycle, with Kenyon cell expression arising only in late pupae (13). Fru^M^ exerts masculinizing effects that could arise at each of these stages. We have begun our analysis here with the adult stage, and observe gene regulatory alterations of synaptic matching molecules, transcription factors, and ion channels that are consistent with neural specificity functions required in the adult. We expect that some of these differences, especially in synaptic matching molecules, would also be observed at earlier developmental stages—synaptic matching molecules are used to guide synapsis and often continue to be expressed to maintain the synapse (55,66,67). At earlier stages, we might begin to observe Fru^M^-dependent regulation of genes required for axon guidance or the induction or suppression of apoptosis. Alternatively, as differentiation is thought to proceed through loss of gene regulatory potential over time, the adult state of these cells as measured by ATAC-seq could represent a summation of all the gene regulatory alterations that occurred during earlier stages of cellular development; this is particularly likely if Fru^M^ decommissions regulatory elements. Finally, we observe in adults that Fru^M^-dependent regulatory elements are cell-type-specific. We cannot rule out the possibility that these or other regulatory elements are used more broadly across *fruitless* neuron types at earlier developmental stages.

### Effects of Fru^M^ on connectivity are unlikely to occur through convergent gene expression

The construction of neural circuits from diverse cellular parts is an extraordinarily complex problem, and the idea that circuit construction could be simplified by expression of the same factors across cells of a circuit has arisen repeatedly as a potentially simplifying mechanism (68–70). Sexually dimorphic transcription factors are particularly compelling examples in which expression of a particular gene appears to “paint” multi-layered circuits dedicated to particular behaviors (1,28,71). Our results here suggest that even when the same transcription factor does label lineally diverse, connected cells, this pattern is unlikely to make the process of establishing circuit connectivity any simpler. First, we suggest that lineally diverse neurons use distinct gene expression programs to activate the shared transcription factor, i.e. that expression of *fruitless* occurs through multiple convergent mechanisms rather than a single mechanism. The extreme length and low exon/intron ratio of many genes that function as neural specificity factors suggest that, like the *fruitless* locus, they are packed with regulatory elements that modularly govern their expression across distinct neuronal populations.

Second, once Fru^M^ is produced, it does not homogenize the expression profiles of these diverse cells, but rather alters expression of distinct gene batteries in each population of cells. While Fruitless is thus shared across these cells, the gene regulatory elements upstream and downstream of it are not. If anything, maintaining Fruitless as a shared node across these cells imposes an additional layer of complexity and constraint on the gene regulatory events that construct the brain. We expect that common expression of transcription factors across layers of sexually dimorphic neural circuits is evolutionarily retained to allow categorization of a set of neuronal transformations as sex-related—these are “switch genes,” not “terminal selectors” (28,72). While examples of shared expression of transcription factors or homophilic adhesion molecules across connected cells will certainly occur from time to time, we do not expect this to be a general model for the construction of circuits.

### Modular control of gene expression

Our findings suggest that aside from *fruitless* itself, there are not genes whose expression is specific to or universal across the circuit. In this view, Fru^M^ does not transcriptionally unify cells of the circuit or simplify circuit specification. Rather, Fru^M^ flags these cells as “male,” and induces male-specific circuit function by tweaking expression of the same effector genes as are used in other combinations in other circuits. The data presented here are consistent with a model where Fru^M^ acts in concert with the distinct transcriptional milieu of each subpopulation of *fruitless* neurons to enact distinct gene regulatory programs and thus alters each class of neurons in unique ways. Gene regulatory programs therefore diverge downstream of Fru^M^. We propose that this modular organization allows evolutionary diversification, as mutations to regulatory elements would be expected to alter gene expression only in individual populations of cells, rather than across the circuit.

## Acknowledgements

We are grateful to the Bloomington Drosophila Stock Center and others who provided flies, as listed in the Resources Table. We thank Nipun Basrur, Patricia Wittkopp, Cameron Palmer, and members of the Clowney lab for critical reading of the manuscript. MVB was supported by the NIH Cellular and Molecular Biology Training Grant T32-GM007315. RD and EJC were supported by fellowships from the Helen Hay Whitney Foundation. EJC is an Alfred P Sloan Research Fellow in Neuroscience and the Rita Allen Foundation Milton Cassell Scholar. Funding was also provided by start-up funds from the University of Michigan.

## Methods

### Resources Table

| Reagent type (species) or resource | Designation | Source or reference | Identifiers | Additional information |
| --- | --- | --- | --- | --- |
| Gene (*Drosophila melanogaster*) | fru |  | FBgn0004652 |  |
| Genetic reagent (*Drosophila melanogaster*) | FruP1-GAL4 (III) | Gift of Barry Dickson | BDSC 66696 | (10) |
| Genetic reagent (*Drosophila melanogaster*) | UAS-mCD8::GFP (II) | Bloomington *Drosophila* Stock Center |  | (73) |
| Genetic reagent (*Drosophila melanogaster*) | MB247-GAL80 (II) | Bloomington *Drosophila* Stock Center | BDSC 64306 |  |
| Genetic reagent (*Drosophila melanogaster*) | FruP1-LexA (III) | Bloomington *Drosophila* Stock Center | BDSC 66698 | (51) |
| Genetic reagent (*Drosophila melanogaster*) | LexAop-tdTomato.Myr (su(Hw)attP5) | César Mendes, Columbia University | FBti0160868 |  |
| Genetic reagent (*Drosophila melanogaster*) | 10XUAS-IVS-myr::GFP {attP2} | Bloomington *Drosophila* Stock Center | BDSC 32197 | (74) |
| Genetic reagent (*Drosophila melanogaster*) | G w[1118]; P{y[+t7.7] w[+mC]=GMR80B02-GAL4}attP2 | Bloomington *Drosophila* Stock Center | BDSC 40064 | (50) |
| Genetic reagent (*Drosophila melanogaster*) | w[1118]; P{y[+t7.7] w[+mC]=GMR84D12-GAL4}attP2 | Bloomington *Drosophila* Stock Center | BDSC 40394 | (50) |
| Genetic reagent (*Drosophila melanogaster*) | w[1118]; P{y[+t7.7] w[+mC]=GMR89D01-GAL4}attP2 | Bloomington *Drosophila* Stock Center | BDSC 46880 | (50) |
| Genetic reagent (*Drosophila melanogaster*) | w[1118]; P{y[+t7.7] w[+mC]=GMR91A09-GAL4}attP2 | Bloomington *Drosophila* Stock Center | BDSC 40571 | (50) |
| Genetic reagent (Drosophila melanogaster) | w[1118]; P{y[+t7.7] w[+mC]=GMR92B12-GAL4}attP2 | Bloomington Drosophila Stock Center | BDSC 48415 | (50) |
| Genetic reagent (Drosophila melanogaster) | w[1118]; P{y[+t7.7] w[+mC]=GMR75F01-GAL4}attP2 | Bloomington Drosophila Stock Center | BDSC 41304 | (50) |
| Genetic reagent (Drosophila melanogaster) | w[1118]; P{y[+t7.7] w[+mC]=GMR80F10-GAL4}attP2 | Bloomington Drosophila Stock Center | BDSC 47070 | (50) |
| Genetic reagent (Drosophila melanogaster) | w[1118]; P{y[+t7.7] w[+mC]=GMR64C09-GAL4}attP2 | Bloomington Drosophila Stock Center | BDSC 39300 | (50) |
| Genetic reagent (Drosophila melanogaster) | P{VT043653-GAL4}attP2 | VDRC | V204214 | (47) |
| Genetic reagent (Drosophila melanogaster) | P{VT043690-GAL4}attP2 | VDRC | V201280 | (47) |
| Genetic reagent (Drosophila melanogaster) | P{VT043692-GAL4}attP2 | VDRC | FBti0169957 | (47) |
| Genetic reagent (Drosophila melanogaster) | P{VT043701-GAL4}attP2 | VDRC | FBti0170490 | (47) |
| Genetic reagent (Drosophila melanogaster) | LexAopFLP (II) | Via Leslie Vosshall | FBal0295487 | (75) |
| Genetic reagent (Drosophila melanogaster) | Tub>gal80> (X) | Gift from Kristin Scott | BDSC 38879 | (76) |
| Antibody | GFP (chicken polyclonal) | Gift from Dawen Cai | n/a | (1:5000) |
| Antibody | dsRed | Takara/Clontech | 632496 | (1:1000) |
| Antibody | nc82 | DSHB | nc82 | (1:25-1:40) |
| Antibody | Anti-chicken Alexa 488 | Fisher | A-11039 | (1:500) |
| Antibody | Anti-rabbit Alexa 568 | Fisher | A-11036 | (1:500) |
| Antibody | Anti-mouse Alexa 647 | Fisher | A-21236 | (1:500) |
|  | DAPI | Sigma | D9542 | 50ng/mL (sorting)<br>1µg/mL (staining) |
|  | 10x PBS, no Ca<sup>2+</sup>, no Mg<sup>2+</sup> | Gibco | 70011-044 |  |
|  | BSA | Sigma | A9085 |  |
|  | Collagenase | Sigma | C0130 | 2mg/mL |
|  | Schneider's Medium | Sigma | S0146 | Sigma Schneider's more reliable than Gibco |
|  | Schneider's Medium | Gibco | 21720024 |  |
|  | 16% PFA | EMS | 15710 | 1% |
|  | Trizol-LS | Fisher | 10296010 |  |
|  | Arcturus Picopure Kit |  | KIT0204 |  |
|  | RNAse free DNase set | Qiagen | 79254 |  |
|  | Zymo-5 columns |  | D4013 |  |
|  | Igepal CA-630 | Sigma | I8896-50mL |  |
|  | Tungsten Wire | California Fine Wire Co | MO285420 |  |
|  | Sylgard 184 | Fisher | 50-366-794 |  |
|  | TD Buffer | Illumina | FC-121-1030 | From Nextera DNA Sequencing Kit |
|  | Tn5 | Illumina | FC-121-1030 | From Nextera DNA Sequencing Kit |
|  | Triton | Sigma | X100 |  |
|  | Normal Goat Serum, lyophilized | MP Biomedical | 8642921 |  |
|  | Fly food | Lab Express | R |  |
|  | Fly food | Lab Express | B |  |
| Software | Bowtie2 |  |  | (77) |
| Software | MACS2 |  |  | (36) |
| Software | deepTools |  |  | (78) |
| Software | DiffBind |  |  | (40,41) |
| Software | i-cisTarget |  |  | (42) |
| Software | HiSat2 |  |  | (80) |
| Software | DESeq2 |  |  | (52) |
| Software | Galaxy |  |  | (81) |
| Software | ChIPSeeker |  |  | (82) |
| Software | DAVID |  |  | (83) |
| Software | gProfiler |  |  | (84) |
| Software | FIMO |  |  | (58) |
| Dataset | FlyFactorSurvey | Fru |  | (85) |
| Dataset | R57C10 neuron ChIP | GSE37032<br>SRP012052 |  | (46) |
| Dataset | roX2 CHART |  |  | (38) |
| Dataset | FruA, FruB, FruC DamID |  |  | (56) |
| Dataset | FruM BirA |  |  | (61) |
| Dataset | Fru neuron TRAP |  |  | (65) |
| Dataset | Fru<sup>Δ</sup>COM ChIP | ENCODE<br>ENCGM860XOW |  | (60) |

### Flies

Flies were maintained on cornmeal-molasses food, or on cornmeal food with a yeast sprinkle (‘R’ or ‘B’ recipes, Lab Express, Ann Arbor, MI) in a humidified incubator at 25C on a 12:12 light:dark cycle. Flies analyzed in all experiments were 2-7 day old adults who were housed in mixed-sex groups.

Genotypes

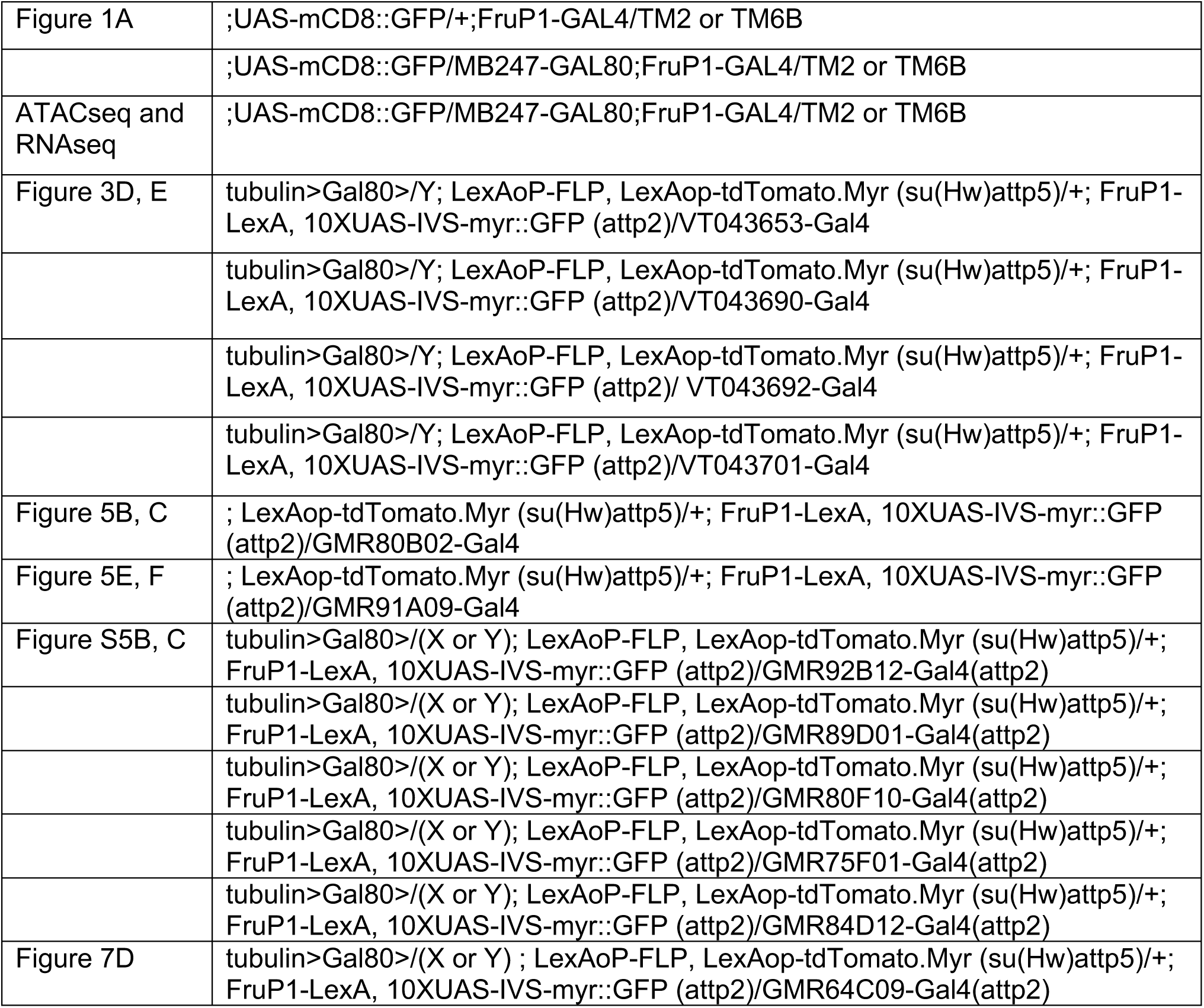

### Flow Cytometry

Brains were dissected for up to 90 minutes in Schneider’s medium supplemented with 1% BSA and placed on ice. Optic lobes were removed during dissection. Brain dissections were interspersed such that both male and female brains were dissected throughout the dissection period. About 20 brains were obtained for each sex. After all dissections were completed, collagenase was added to a final concentration of 2mg/mL and samples were incubated at 37C for 20 minutes, without agitation.

Samples were dissociated by trituration and spun down at 300g, 4C, for 5 minutes. Collagenase solution was removed and replaced with PBS+0.1% BSA, and cells were passed through a cell strainer cap and supplemented with 50ng/mL DAPI before being subjected to flow cytometry on an a FACS Aria II. Plasticware for cell dissociation and collection was pre-treated by rinsing with PBS+1% BSA to prevent cells from sticking to bare plastic.

During flow cytometry, dead and dying cells were excluded using DAPI signal, and forward scatter and side scatter measurements were used to gate single cells. Using our dissociation methods, 50-80% of singlets appeared viable (DAPI-low), and 2-5% of viable singlets were GFP^+^. We collected 6,000-10,000 GFP^+^ cells for each *fru*^+^ sample and analyzed matched numbers of GFP^-^/*fru*^-^ cells. For each replicate, we sorted male and female cells during the same session and performed transposition or RNA extraction in parallel. During sorting, we made two adjustments to protect the fly primary cells, which were very delicate—we disabled agitation of the sample tube, and sorted using the “large nozzle,” e.g. 100μm, i.e. using larger droplet size and lower pressure. For ATAC-seq, we sorted cells into PBS supplemented with 0.1% BSA. For RNA-seq, we sorted cells directly into Trizol-LS.

### Brain dissections, staining, and imaging

Brains were dissected in external saline (108 mM NaCl, 5 mM KCl, 2 mM CaCl2, 8.2 mM MgCl2, 4 mM NaHCO3, 1 mM NaH2PO4, 5 mM trehalose, 10 mM sucrose, 5 mM HEPES pH7.5, osmolarity adjusted to 265 mOsm). For two-photon imaging, brains were then transferred fresh to 35mm imaging dishes and pinned to sylgard squares with tungsten wire. Imaging was performed on a Bruker Investigator using a 1.0 NA, 20x, water-dipping objective. Stacks were collected along the anterior-posterior axis with 1 micrometer spacing in Z and ∼350nm axial pixel size.

For immunostaining and confocal imaging, brains were dissected for up to twenty minutes before being transferred to 1% paraformaldehyde in PBS, on ice. All steps were performed in cell strainer baskets (caps of FACS tubes) in 24 well plates, with the brains in the baskets lifted from well to well to change solutions. Brains were fixed overnight at 4C in 1% PFA in PBS. On day 2, brains were washed 3×10’ in PBS supplemented with 0.1% triton-x-100 on a shaker at room temperature, blocked 1 hour in PBS, 0.1% triton, 4% Normal Goat Serum, and then incubated for at least two overnights in primary antibody solution, diluted in PBS, 0.1% triton, 4% Normal Goat Serum. Primary antibody was washed 3×10’ in PBS supplemented with 0.1% triton-x-100 on a shaker at room temperature, then brains were incubated in secondary antibodies for at least two overnights, diluted in PBS, 0.1% triton, 4% Normal Goat Serum. DAPI (1 microgram/mL) was included in secondary antibody mixes. Antibodies and concentrations can be found in the resources table.

Brains were mounted in 1x PBS, 90% glycerol supplemented with propyl gallate in binder reinforcement stickers sandwiched between two coverslips. Samples were stored at 4C in the dark prior to imaging. The coverslip sandwiches were taped to slides, allowing us to perform confocal imaging on one side of the brain and then flip over the sandwich to allow a clear view of the other side of the brain. Scanning confocal stacks were collected along the anterior-posterior axis on a Leica SP8 with 1 micrometer spacing in Z and ∼150nm axial pixel size.

### Assay for Transposase-Accessible Chromatin

After FAC-sorting, nuclear isolation and Tn5 transposition were performed as in (29) with modifications made for small numbers of cells as described in (35). Transposed DNA was isolated and stored at −20C until library preparation. Transposed DNA was amplified with barcoded primers in NEBNext High Fidelity 2X PCR Master Mix (NEB) and purified with Ampure XP beads (Beckman Coulter) at a ratio of 1.6uL beads per 1uL library. Purified library was eluted in 30uL of 10mM Tris-HCl pH8, 0.1mM EDTA.

The quality of prepared libraries was verified with a Bioanalyzer 2100 using a high sensitivity DNA kit (Agilent). Libraries were quantified using KAPA qPCR assay (KAPA Biosystems), and multiplexed and sequenced on a NextSeq Illumina machine with 75 base pair, paired-end reads. Libraries were sequenced to a depth of 35-70 million reads.

#### Processing of ATAC-seq data

Adapters were trimmed using cutadapt and reads below 18 bases were discarded. Reads were aligned to dm6 with Bowtie2 with option –X 1000 to set the maximum fragment length for paired-end reads. About 70% of reads mapped uniquely to the fly genome. Aligned reads were processed with samtools to create a bam file. Picard MarkDuplicates was used to mark duplicates and samtools was used to generate a final bam file with de-duplicated reads passing a q30 quality filter. After de-duplication, we obtained 8.9-16.3 million unique reads per library.

The sequencing data were uploaded to the Galaxy web platform, and we used the public server at usegalaxy.org to analyze the data (81). Peaks were called with MACS2 “callpeak” using – nomodel at an FDR threshold of <0.001, which gave a reproducible and robust peak calls across replicates and by eye.

#### Analysis of Differential Accessibility

For comparisons between neurons of the same sex status (i.e. male fru+ vs male fru- or female fru+ vs female fru-), DiffBind was run with default parameters using filtered bam files for each replicate and MACS2 called peaks for each replicate as input.

In female flies, the X chromosomes accounted for 20-21% of all ATAC-seq reads, while only accounted for 14% of all male ATAC-seq reads across samples. For comparisons of chromatin accessibility between sexes, this difference in X:A DNA ratio introduced extremely large bias in regions called as differentially accessible. We initially identified 1374 differentially accessible regions between male fru-neurons and female fru-neurons, compared to the 95 regions reported in this paper. This is due to whole-genome rather than per-chromosome normalization performed by DiffBind and results in artificially high numbers of regions to be female-biased on X and male-biased on the autosomes.

Therefore, we separated aligned reads from the X chromosome and autosomes using “Slice BAM by genomic regions”, selecting for chrX, or for chr2L, 2R, 3L, 3R, and 4. DiffBind was run using the same MACS2 called peaks, but with either X chromosome or autosome aligned reads. Separate lists of changed regions (FDR <0.05) were concatenated per comparative analysis.

Tables of genomic regions were downloaded and processed in R as follows: Peaks were annotated using ChIPSeeker – nearest gene, +/-50bp promoter and exported as tables. Fru^M^-specific regions with annotated protein-coding genes were filtered for unique genes. List of genes were loaded into gProfiler with default settings. Annotation from combined MACS2 peak calls used as custom background list. Fru^M^-specific gene lists were also loaded into DAVID for domain annotation, without a custom background.

### RNA sequencing

#### RNA extraction

Cells were sorted into Trizol-LS and stored at −80C until RNA isolation. For RNA extraction, we followed the standard Trizol-LS protocol until the aqueous phase was isolated. We then passed the aqueous phase over Arcturus Picopure columns, including a DNAse treatment on the column. Our protocol is copied from the following document, with our thanks to the authors, who are unknown to us. We include an image of the document here in case the link becomes inactive in the future:

https://wiki.library.ucsf.edu/download/attachments/188645378/RNA%20purification%20from%20Trizol%20samples%20via%20PicoPure%20column.pdf?version=1&modificationDate=1349799100000&api=v2

RNA purification from Trizol samples using Picopure columns Ed R. 20101207

ABI Arcturus Picopure Kit #

Qiagen RNAse-free DNAse set #79254

Modifications from ABI help line for Arcus Picopure (Candice) 12/7/10

1. Trizol sample:
  a. Process with 0.2 vol of chloroform
  b. Spin max speed for 5 minutes (start picopure column conditioning)
  c. Take aqueous layer
  d. Add equal volume of 70% Ethanol (RNAse free)
2. PicoPure column conditioning
  a. Condition picopure column for 5 minutes
  b. Spin out at max speed for 1 minute
3. Bind RNA to picoPure column
  a. Load up to 300 ul of the supernatant/70% EtOH mix from 1d at a time
  b. Spin at 100g x 2 minutes for each load.
  c. After last load, spin 16000g x 30 seconds
  d. Discard flowthrough
4. Wash with 100 ul Wash Buffer 1 (WB1) at 8000g x 1 minute
5. DNAse treatment (optional, but a good idea because of trizol)
  a. For each sample, combine 35 ul of RDD buffer with 5 ul of DNAseI stock solution (previously resuspended per Qiagene protocol)
  b. Add 40 ul of DNAse mix onto membrane.
  c. Incubate RT x 15 minutes
  d. Add 40 ul of Wash Buffer 1 (WB1) and spin 8000g x 15 sec
6. Wash with 100 ul of Wash buffer 2 (WB2) at 8000g x 1 minute
  a. Empty flow through
7. Wash again with 100 ul Wash Buffer 2 (WB2) at 16000g x 2 minutes
  a. Check to make sure no wash buffer remains
8. Elution step
  a. Transfer to a new 0.5 ml tube (in kit)
  b. Add 11 ul of 42C pre-warmed elution buffer to membrane
  c. Wait 1 minute
  d. Spin at 1000g x 1 minute to distribute buffer
  e. Elute at 16000g x 1 minute
  f. Consider doing a 2^nd^ elution to capture as much flowthrough as possible
9. Store at −80C.

General notes

Using the Qiagen RNeasy columns → significant loss of > 50%

RNA quantification and quality assessment were performed on an RNA TapeStation at the UMich Genomics Core. We typically obtain 0.1-0.3pg of RNA per cell, depending on cell type and developmental stage. Note that insect 28S rRNA is processed to a size similar to 18S rRNA, thus “RNA Integrity Number” or similar that are calculated by these machines will not reflect the true RNA quality.

#### Library preparation and sequencing

RNA libraries were prepared by the UMich Genomics Core with the following protocol: Samples were subject to quality control on an RNA TapeStation. Total starting RNA was around 1 ng per sample. Library preparation was performed using the NEBNext® Single Cell/Low Input RNA Library Prep Kit for Illumina, with amplification cycles calibrated to the amount of total RNA. Unstranded, poly-A selected libraries were sequenced on an Illumina NovaSeq using 150bp paired end reads to a depth of ∼30 million reads per library.

#### Alignment and analysis of RNA-seq data

All processing steps were done through the Galaxy web platform. Reads were trimmed with trim galore! with automatic adapter detection and aligned to dm6 using HiSAT2 in paired end mode. Uniquely aligning reads above MAPQ 30 were selected using JSON and MarkDuplicates, retaining 25.4-39.6 million reads per library. FastQC and MultiQC were used to visualize quality metrics across samples. Coverage tracks were generated using deeptools “bamCoverage”.

Exon reads were counted and aggregated at a gene-level using featureCounts. TPM values were calculated using StringTie and averaged between replicates. Differential expression was determined using DESeq2 using an p.adj threshold of >0.05 for reporting significance.

### Additional Genomic Analyses

#### Analysis of lncRNA:roX2 tethering near sites of sex-specific chromatin accessibility

Genomic locations of *lncRNA:roX2* binding to chromatin sing CHART was lifted over from dm3 into dm6 using UCSC LiftOver. Lifted-over bed files were sorted using bedtools “sortBed” and relative distance of lncRNA:roX2 tethering sites was determined using bedtools “RelDist” for the X chromosomes only.

#### Neuron-specific histone marks

Histone chromatin immunoprecipitation from adult neurons (46) was downloaded in FASTQ format from SRA (86) using “Download and Extract Reads in FASTQ”, trimmed using Trim Galore! (87) with automatic adapter detection. Trimmed reads were aligned to dm6 using Bowtie2 and made into coverage tracks for comparison using deepTools “bamCoverage”.

#### Fru DamID analysis

Ratio files of FruA, FruB, and FruC DamID enrichment over dam-only control were downloaded in GFF format from GEO (GSE52247). Regions were resized to remove 5bp on each flank, thus removing overlapping regions of microarray probes and rebased to bed/bedGraph compatible format. Genomic regions were then lifted over from dm3 into dm6 using RLiftOver. Tables were exported as bedGraph files, uploaded to the Galaxy web platform and converted from bedGraph to bigWigs using “Wig/BedGraph-to-bigWig” using default parameters.

#### Analysis of Fru motifs

All ATAC-seq genomic regions from DiffBind comparison (FDR >= 1) between female *fru*+ and male *fru*+ were converted to FASTA using bedtools MakeFasta. FASTA sequences were run through the web version of FIMO using Fru motifs identified by SELEX (57) with no threshold (Q val = 1) to re-capture all sequences which were input. We used p-values rather than Q-values in our analysis because the SELEX motifs are 8bp, and near-identical matches to sequences do not stand up well to p-value correction. The top match per region per Fru isoform was selected using R based on lowest p-value match. Regions previously selected (Fig. 4) as Fru^M^-specific were flagged. Tables were converted to cumulative frequencies and plotted.

### Data access

Data associated with this paper will be deposited on GEO prior to publication and made available to reviewers during peer review. Tables of differential accessibility and differential expression across comparisons are provided in supplemental tables 1 and 2. At publication, we plan to release a UCSC trackhub include our datasets and published datasets re-analyzed here.

